# Multi-omics analysis reveals CMTR1 upregulation in cancer and roles in ribosomal protein gene expression and tumor growth

**DOI:** 10.1101/2024.10.30.621171

**Authors:** Ion John Campeanu, Yuanyuan Jiang, Hilda Afisllari, Sijana Dzinic, Lisa Polin, Zeng-Quan Yang

## Abstract

The mRNA cap methyltransferase CMTR1 plays a crucial role in RNA metabolism and gene expression regulation, yet its significance in cancer remains largely unexplored. Here, we present a comprehensive multi-omics analysis of CMTR1 across various human cancers, revealing its widespread upregulation and potential as a therapeutic target. Integrating transcriptomic and proteomic data from a large set of cancer samples, we demonstrate that CMTR1 is upregulated at the mRNA, protein, and phosphoprotein levels across multiple cancer types. Functional studies using CRISPR-mediated knockout and siRNA knockdown in breast cancer models show that CMTR1 depletion significantly inhibits tumor growth both *in vitro* and *in vivo*. Transcriptomic analysis reveals that CMTR1 primarily regulates ribosomal protein genes and other transcripts containing 5’ Terminal Oligopyrimidine (TOP) motifs. Additionally, CMTR1 affects the expression of snoRNA host genes and snoRNAs, suggesting a broader role in RNA metabolism. Mechanistically, we propose that CMTR1’s target specificity is partly determined by mRNA structure, particularly the presence of 5’TOP motifs. Furthermore, we identify a novel CMTR1 inhibitor, N97911, through *in silico* screening and biochemical assays, which demonstrates significant anti-tumor activity *in vitro*. Our findings establish CMTR1 as a key player in cancer biology, regulating critical aspects of RNA metabolism and ribosome biogenesis, and highlight its potential as a therapeutic target across multiple cancer types.

## Introduction

The mRNA cap, a highly methylated modification at the 5’ end of RNA polymerase II (RNA Pol II)-transcribed RNAs, plays multiple critical roles in eukaryotic gene expression [1–7]. It protects RNA from degradation both during transcription and in the cytoplasm, recruits protein complexes involved in RNA processing, nuclear export, and translation initiation, and marks cellular mRNA as ‘self’ to prevent degradation by the innate immune system [1–7]. The formation and methylation of the cap structure are catalyzed by a set of specialized enzymes. During the early stages of transcription, RNGTT (RNA guanylyltransferase and triphosphatase) attaches the inverted guanosine cap to the first transcribed nucleotide via a triphosphate bridge [1, 2, 7, 8]. Subsequently, a series of cap methyltransferases—including RNMT (RNA guanine-7 methyltransferase), CMTR1 (cap methyltransferase 1), CMTR2 (cap methyltransferase 2), and/or PCIF1 (phosphorylated CTD interacting factor 1)—methylate specific sites on the guanosine cap and the first two transcribed nucleotides [1–7]. This process protects and stabilizes mRNA while serving as a crucial regulatory mechanism for gene expression, influencing various cellular processes including development, differentiation, and disease progression.

CMTR1 and CMTR2, which contain a unique Rossmann fold methyltransferase domain known as Ftsj, catalyze 2’-O-methylation of the first and second transcribed nucleotides, respectively [9–14]. However, CMTR1 and CMTR2 contain distinct functional domains flanking their Ftsj methyltransferase domains [12]. Additionally, CMTR1 in mammals, including humans, is predominantly nuclear, while CMTR2 is primarily cytoplasmic [13]. Notably, CMTR1 interacts directly with RNA polymerase II via its WW domain, preferentially binding to Ser-5 phosphorylated C-terminal domains (CTDs) [15, 16]. This interaction allows CMTR1 to be recruited effectively to transcription start sites, correlating with RNA polymerase II abundance. Recent studies also revealed that CMTR1 has gene-specific impacts on transcript abundance and plays a significant role in embryonic stem cell differentiation, particularly in maintaining expression of histone and ribosomal protein genes [17]. As CMTR1 is a known interferon-stimulated gene, it also plays roles in immune-mediated pathways during viral infection [16, 18–20]. CMTR1 inhibition provides strong protection against infection by multiple influenza A strains, and synergizes with the viral endonuclease inhibitor baloxavir, which blocks influenza A infection by preventing cap snatching [18]. Given its fundamental roles in RNA metabolism and gene regulation, it is not surprising that alterations in CMTR1 are linked to various human diseases, including cancer [21–23]. This connection underscores the potential of this enzyme as a novel therapeutic target and biomarker in disease progression and treatment.

In this study, we aimed to understand the genetic and transcriptomic patterns, as well as the clinical significance, of cap methyltransferases—focusing primarily on CMTR1—by employing an unbiased multi-omics approach across a large dataset of various human cancers. We used genetic approaches to inhibit CMTR1 and identified its downstream targets in both human and mouse cancer models. Additionally, we explored *in silico* screening and biochemical assays to identify novel CMTR1 inhibitors. Our results highlight the therapeutic potential of targeting CMTR1 in multiple cancer types.

## Results

### Differential expression and phosphorylation of CMTR1 and CMTR2 across cancer types

Previously, we employed an unbiased approach to investigate genetic alterations in over 50 methyltransferases across human cancers. This analysis led to the identification of FTSJ3, a 2’-O-methyltransferase, as a potential regulator of breast cancer progression [24]. In human cells, two Ftsj-domain-containing methyltransferases, CMTR1 and CMTR2, specifically target the mRNA cap [2, 3, 5–7]. However, our understanding of the expression patterns and biological roles of CMTR1 and CMTR2 in human cancers remains limited.

To address this knowledge gap, we analyzed mRNA expression changes of CMTR1 and CMTR2 in cancerous tissues relative to normal tissues across 15 tumor types from The Cancer Genome Atlas (TCGA) dataset. We selected tumor types with at least ten normal tissue samples available for comparison. Our analysis revealed that CMTR1 exhibited significantly (*p* < 0.05) increased RNA levels in five TCGA cancer types compared to normal adjacent tissues (NATs): breast (BRCA), bladder (BLCA), colorectal (COADREAD), head and neck (HNSC), and liver (LIHC) cancers (Figure 1A). Conversely, CMTR2 showed significantly decreased RNA levels in nine tumor types: BRCA, HNSC, LIHC, kidney clear cell (KIRC), kidney papillary (KIRP), lung adenocarcinoma (LUAD), lung squamous cell carcinoma (LUSC), prostate adenocarcinoma (PRAD), and endometrial carcinoma (UCEC) (Supplementary Figure 1A & Supplementary Table 1).

**Figure 1:**
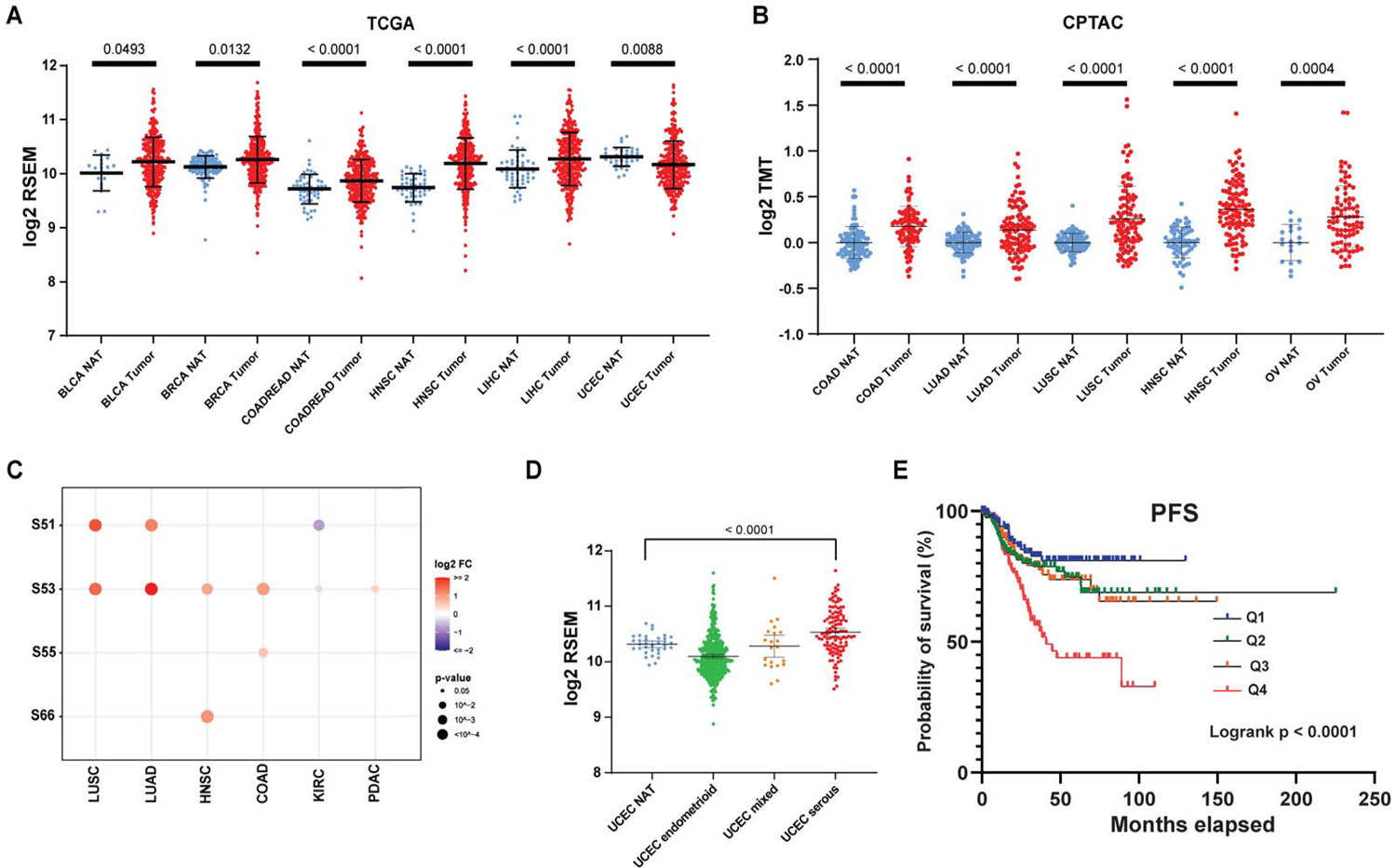
CMTR1 expression, phosphorylation, and prognostic value across multiple cancer types (A) Dot plots of CMTR1 mRNA expression (log2 RSEM) in normal and tumor samples from six TCGA cancer types. (B) Dot plots of CMTR1 protein abundance (log2 TMT) in normal and tumor samples from five CPTAC cancer types. Blue dots represent normal adjacent tissue (NAT) samples, whereas red dots represent tumor samples. Each data point represents an individual patient sample. The p-value from Mann-Whitney U test comparing tumor and NAT samples for each cancer type is displayed. Error bars indicate standard deviation of mRNA or protein expression levels across patients. (C) Normalized phosphorylation levels of CMTR1 residues S51, S53, S55, S63, and S66 relative to total CMTR1 protein abundance in normal and tumor samples from six CPTAC cancer types. The heatmap displays log2 FC for cancer types with at least 10 samples in both tumor and NAT groups. The color gradient from purple to red represents the degree of log2 FC between tumor and normal samples. Dot size indicates statistical significance. Only data points with log FC >|0.2| and *p*-value < 0.05 are shown. (D) mRNA expression levels of CMTR1 in TCGA uterine corpus endometrial carcinoma (UCEC) by histological subtypes. CMTR1 expression is significantly higher in the serous subtype of UCEC compared to normal adjacent tissue (NAT) samples, while it is lower in the endometrioid subtype compared to NAT samples. (E) Kaplan-Meier progression-free survival (PFS) curves of TCGA UCEC patients (n = 530) stratified into quartiles (Q1, Q2, Q3 and Q4) by CMTR1 expression. The highest expression group (Q4) is shown with a red line.

The recent availability of proteomic profiles across a broad range of cancers from Clinical Proteomic Tumor Analysis Consortium (CPTAC) projects has provided an unprecedented opportunity to investigate proteomic changes of CMTRs in cancers relative to normal tissues [25]. We analyzed protein expression differences between tumors and NATs using CPTAC proteomics data for eight tumor types, each with at least ten NAT samples available. Our analysis revealed that CMTR1 protein was significantly upregulated in five tumor types compared to their respective NAT samples: ovarian (OV), colon (COAD), lung squamous cell carcinoma (LUSC), HNSC, and LUAD (Figure 1B). Interestingly, we found that CMTR2 protein was significantly upregulated in several tumor types, including HNSC, KIRC, LUAD, and LUSC (Supplementary Figure 1 & Supplementary Table 1). This upregulation occurred even in cases where CMTR2 mRNA levels were downregulated, such as in HNSC or LUSC, compared to NATs (Supplementary Figure 1). These results suggest an additional layer of regulation for CMTR2 protein expression, independent of mRNA expression, in certain tumor types.

Given that phosphorylation is a key process in regulating protein activity, we also analyzed CPTAC phosphoproteomics data to compare changes in CMTR1 phosphorylation between tumor and normal samples [26]. Based on the PhosphoSitePlus database, we found that CMTR1 phosphorylation sites are enriched in the N-terminal region of the protein, while CMTR2 lacks this phosphorylation-enriched region (data not shown) [27]. Additionally, phosphorylation sites of CMTR2 were barely detected in the CPTAC samples. Therefore, we focused on CMTR1 phosphorylation sites and levels in CPTAC samples. Three phosphorylation sites—S51, S53, and S66—were detected in at least five of eight CPTAC tumor types (with at least ten samples in both tumor and normal groups), allowing for comprehensive analysis. Comparing phosphorylation levels between CPTAC tumor and normal samples, we found that phosphorylation levels of S51 and S53 were significantly upregulated in four CPTAC tumor types (Supplementary Figure 2, Supplementary Table 2). After normalizing phosphorylation levels to CMTR1 protein abundance we observed similar results, with S53 phosphorylation levels upregulated in five CPTAC tumor types (Figure 1C) [28].

Our analysis of CMTR1 data revealed an unexpected reduction in median RNA levels in all TCGA endometrial cancer (UCEC) samples, contrasting with its expression patterns in other tumor types (Figure 1A). To further investigate this finding, we stratified UCEC tumors by histological subtype (endometrioid, serous, and mixed) [29–31]. This analysis showed that CMTR1 expression is either decreased (at the RNA level) or unchanged (at the protein level) in UCEC’s endometrioid subtype (Figure 1D, Supplementary Figure 3A). Conversely, UCEC’s serous subtype, which is more aggressive than other subtypes, exhibited increased CMTR1 expression at both RNA and protein levels (Figure 1D, Supplementary Figure 3A). Importantly, we found that increased CMTR1 expression correlates with reduced overall and progression-free survival in TCGA UCEC tumor samples (Figure 1E, Supplementary Figure 3B). This effect appears to be largely driven by the increased expression in serous subtype tumors, which are known to have poor prognoses in UCEC [32]. Furthermore, when grouping CMTR1 protein expression in CPTAC data by molecular subtype, we also observed increased expression in copy number variation (CNV)-high tumors compared to both normal tissue and other UCEC molecular subtypes (Supplementary Figure 3C). The CNV-high molecular subtype is mainly comprised of tumors with serous histology [31].

In conclusion, our comprehensive analysis revealed that 2’-O capping enzymes, particularly CMTR1, undergo significant alterations across multiple cancer types at the RNA, protein, and phosphorylation levels.

### *In vitro* and *in vivo* evidence for CMTR1’s tumor-promoting function

To investigate the biological importance of CMTR1 and CMTR2 in various tumors, we analyzed genome-wide CRISPR screen data (DepMap 24Q2) from more than 1,000 tumor cell lines [33, 34]. CMTR1 exhibited significantly lower cancer dependency scores compared to CMTR2, with mean scores of −0.79 and −0.08, respectively (p < 0.0001). Lower scores indicate greater criticality for tumor cell growth and survival. Notably, 19.3% of cancer cell lines showed CMTR1 scores at or below −1 (the average score of critical cell survival genes), whereas only 0.09% of cell lines exhibited such scores for CMTR2 (Supplementary Figure 4A). To further evaluate CMTR1’s role in cancer cells, we employed CRISPR gene editing to knockout (KO) CMTR1 in the 4T1 breast cancer cell line. We selected the 4T1 line due to its molecular similarity to human basal-like breast cancer, which also shows relatively high CMTR1 expression in CPTAC breast cancers (Supplementary Figure 4B). In addition, our previous study of more than 50 RNA methyltransferases in TCGA breast cancers revealed that mRNA levels of CMTR1, but not CMTR2, are significantly higher in the basal-like subtype compared to the ER-positive luminal subtype [24]. Analysis of CMTR1 levels in CCLE breast cancer cell lines grouped by subtype also showed increased expression in basal-like cell lines (Supplementary Figure 5A). Finally, Western blotting assay of CMTR1 protein levels in breast cancer lines revealed that protein expression of CMTR1 was dramatically higher in a subset of basal-like lines, including COLO824 and MDA-MB-468 (Supplementary Figure 5B). CRISPR-mediated editing of CMTR1 in 4T1 cells was confirmed by DNA sequencing, and CMTR1 depletion was validated by Western blot assay (Figure 2A). CMTR1 depletion dramatically inhibited 4T1 cell growth compared to negative control cells (Figure 2B). Clonal growth efficiency in soft agar is a measure of the ability of cancer cells to form colonies in a semi-solid medium, which is considered a hallmark of tumorigenicity [35]. We found that CMTR1 KO significantly reduced 4T1 anchorage-independent growth in soft agar (Figure 2C). Invasion assays showed that CMTR1 KO significantly inhibited the invasive capacity of 4T1 cells (Figure 2D, Supplementary Figure 6). To corroborate these findings in human cancer cells, we used siRNA to knockdown CMTR1 in MDA-MB-436 human basal-like breast cancer cells. qRT-PCR and western blot assays revealed that siRNA significantly decreased the expression of CMTR1 at both mRNA and protein levels in MDA-MB-436 cells (Figure 2E). Consistent with our observations in 4T1 cells, CMTR1 knockdown in MDA-MB-436 cells also inhibited growth *in vitro* (Figure 2F).

**Figure 2:**
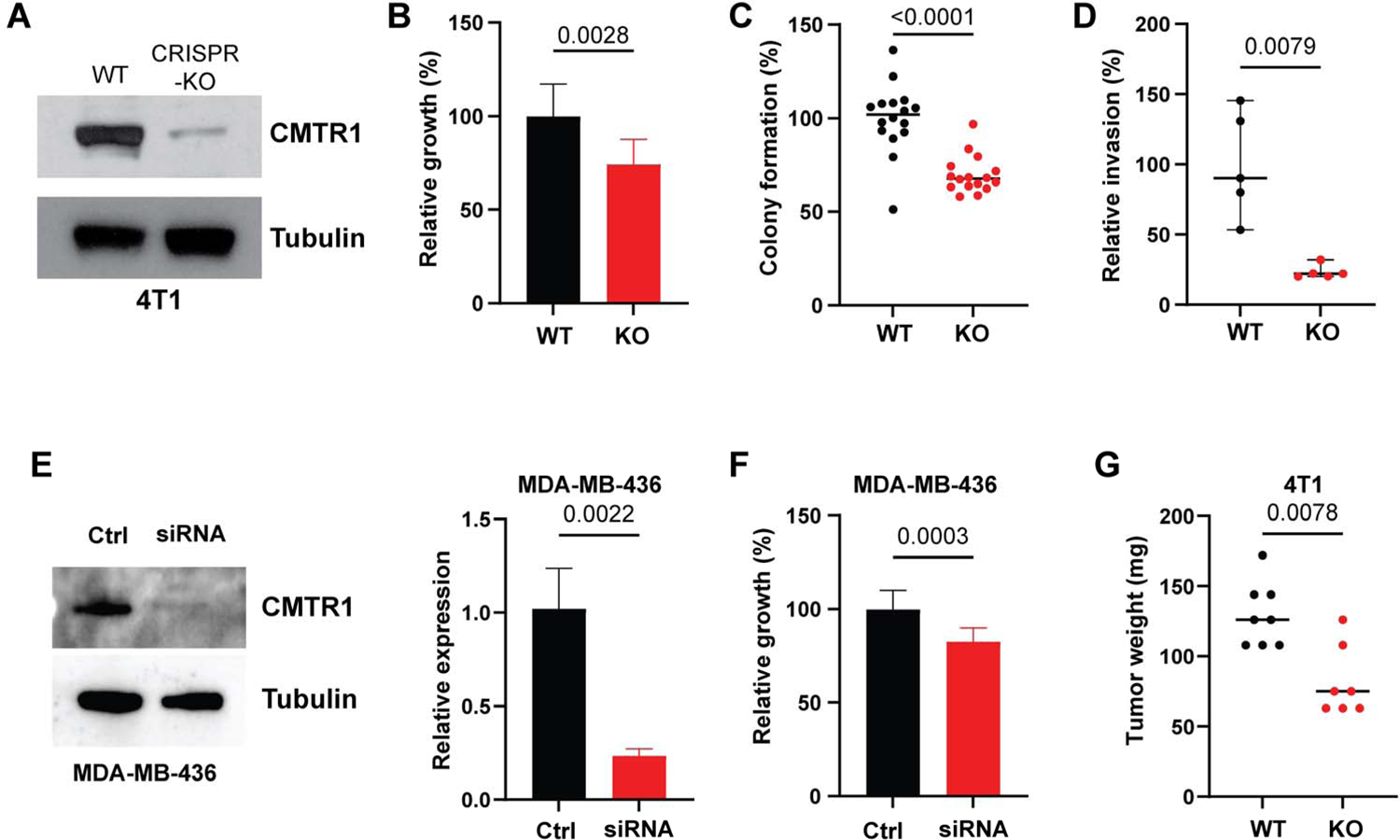
CMTR1 reduces tumor growth and invasion *in vitro* and *in vivo*. (A) Immunoblot showing the expression of CMTR1 in the CMTR1-knockout (KO) 4T1 cells. (B) *in vitro* growth, (C) colony formation in soft agar, and (D) invasion in CMTR1-KO and control 4T1 cells. (E) Knockdown of CMTR1 in human basal-like breast cancer MDA-MB-436 cells with the siRNA was confirmed by qRT-PCR and western blot assays. (F) Bar graph shows relative cell growth after knocking down CMTR1 in MDA-MB-436 cells. (G) Tumor weight of *in vivo* 4T1 tumors with or without CMTR1 knockout. Error bars: SD.

To assess the *in vivo* relevance of our findings, we injected 4T1 CMTR1-KO cells into mice. CMTR1 depletion significantly inhibited tumor growth *in vivo* (Figure 2G). In summary, our data strongly suggests that CMTR1 plays a crucial tumor growth-supporting role across multiple cell types and experimental models.

### CMTR1 depletion leads to downregulation of ribosomal protein genes across cancer cell lines

We next investigated the transcriptomic effects of CMTR1 depletion using our 4T1 CRISPR KO model. RNA-Seq analysis revealed 196 genes with reduced expression and 21 genes with increased expression in CMTR1 KO cells, using a stringent cut-off for our RNA-seq analysis [adjusted p-value (q-value) < 0.05] (Figure 3A, Supplementary Table 3A). Gene Ontology (GO) analysis showed that downregulated genes were significantly associated with two pathways: ribosome (*p* = 5.20E-74, *q* = 2.34E-72) and oxidative phosphorylation (*p* = 0.0019, *q* = 0.04577). Notably, 57 out of 196 downregulated genes were ribosomal pathway members (Figure 3A, Supplementary Table 3B). No significant pathways were identified for upregulated genes. To validate these findings, we analyzed published RNA-Seq datasets from the lung cancer cell line A549 with or without CMTR1 depletion, previously used to identify host dependency factors for influenza A virus infection [18]. In A549 cells, with and without interferon (IFN) treatment (A549NT and A549IFN respectively), we confirmed that the top downregulated genes were significantly enriched in ribosome pathways, primarily ribosomal proteins (*p* = 2.82E-55, *q* = 8.73E-53 and *p* = 6.59E-46, *q* = 2.07E-43, respectively) (Supplementary Figure 7, Supplementary Tables 4-5). Intersecting all three datasets (4T1, A549NT, and A549 IFN) revealed 64 genes commonly downregulated, while only one gene, LARP4 (La ribonucleoprotein 4), was commonly upregulated (Figure 3B, Supplementary Table 6A). GO analysis of the common downregulated gene set showed 53 of 64 genes were ribosomal protein (RP) genes (Figure 3B, Supplementary Table 6B). We confirmed the downregulation of multiple RP genes (e.g., RPS10, RPS14, and RPL13A) upon CMTR1 loss via qRT-PCR in both our 4T1 KO model and an MDA-MB-231 CMTR1 siRNA model (Figure 3C, Supplementary Figure 8).

**Figure 3:**
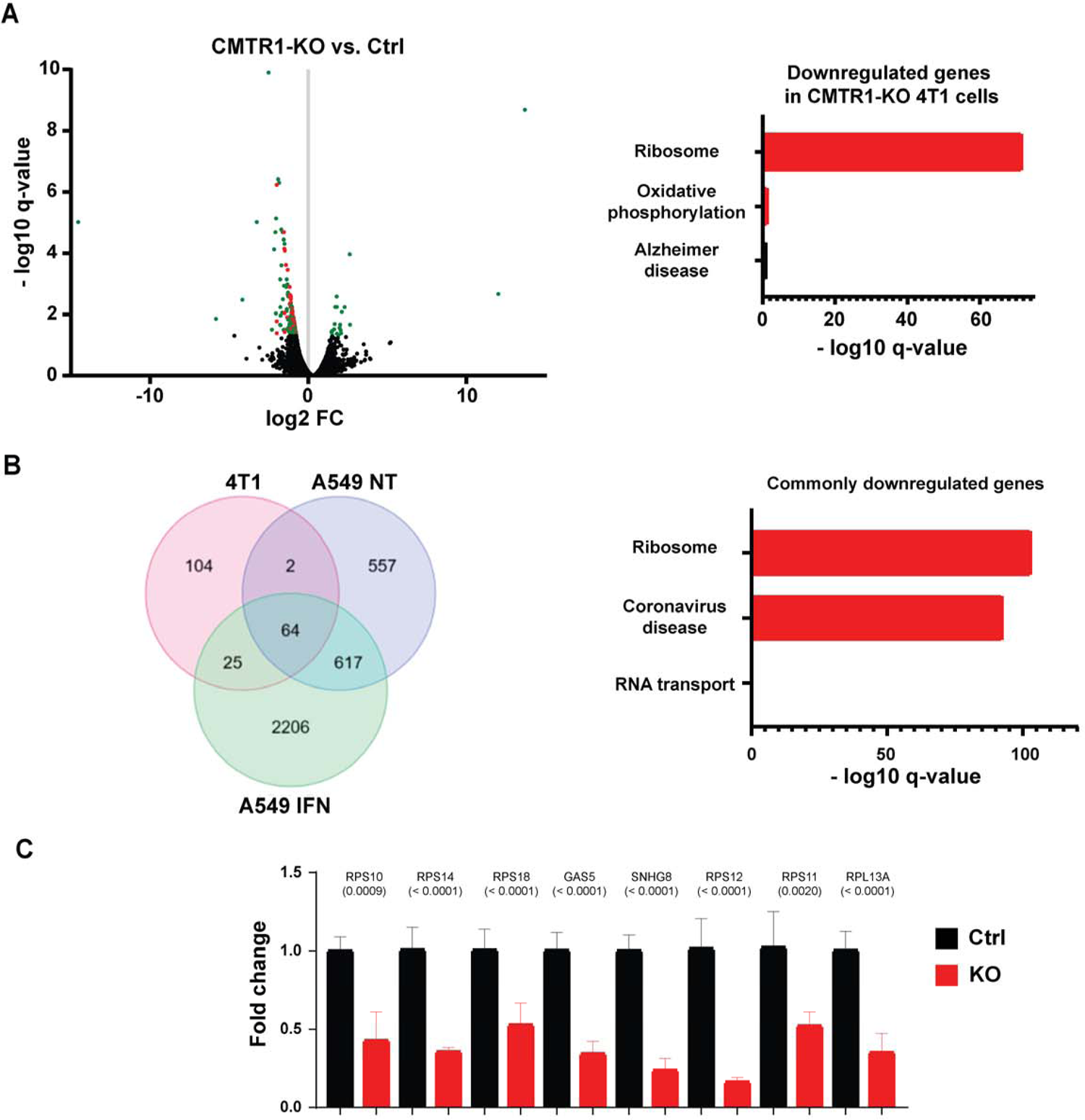
CMTR1 loss reduces ribosomal protein gene expression in cancer cell lines. (A) Volcano plot of differentially expressed genes in 4T1 cells after CMTR1 knockout (left), with pathway analysis of downregulated genes (right). Each dot represents a gene. Green dots indicate significantly (*q* < 0.05) up-and down-regulated genes in CMTR1 knockout 4T1 cells, while red dots highlight significantly down-regulated ribosomal protein genes or SNHGs (snoRNA non-coding host genes). In the right panel, Enrichr pathway analysis using the KEGG 2019 Mouse library was performed on the significantly down-regulated genes in CMTR1 knockout 4T1 cells. For the pathway analysis, red is significant, and the longer the bar the more significant the pathway. (B) Intersection of genes downregulated upon CMTR1 knockout (KO) in three cell models (4T1, A549 untreated, and A549 IFN-treated; left), and pathway analysis of commonly downregulated genes (right). In the right panel, Enrichr pathway analysis using the KEGG 2021 Human library was performed on the common significantly down-regulated genes in CMTR1 knockout cells. For the pathway analysis, red is significant, and the longer the bar the more significant the pathway. (C) Relative expression levels of eight ribosomal protein genes or non-coding SNHGs measured by qRT-PCR in 4T1 cells after CMTR1 knockout. *p*-values for each gene, determined by Welch’s t-test, are displayed. Ctrl: Control; KO: CMTR1 knockout. Error bars: SD.

In summary, our data, together with previously published findings, strongly indicate that ribosomal protein genes are among the primary targets regulated by CMTR1 across various cancer types.

### Enrichment of TOP elements in CMTR1-regulated genes

Expression of ribosomal proteins (RPs) and other factors required for protein synthesis is crucial for cellular function. The 5’ Terminal Oligopyrimidine (TOP) motif, characterized by a cytosine as the first nucleotide followed by 4-15 pyrimidines (C/U), is found in transcripts encoding all human RPs, as well as various initiation factors, elongation factors, and other proteins essential for translation [36]. Given that RP genes were among the most significantly modulated transcripts following CMTR1 depletion in cancer cells, we hypothesized that CMTR1-regulated transcripts are enriched for 5’ TOP elements.

To test this hypothesis, we utilized a recently developed TOP score metric tool to quantify TOP elements in transcripts [37]. We calculated TOP scores from our 4T1 transcription profiles following CMTR1 depletion, as well as the common gene set. Our analysis revealed that genes downregulated after CMTR1 depletion were significantly enriched for TOP elements, exhibiting dramatically increased TOP scores compared to other genes (Figure 4A). These data suggest that CMTR1 deficiency predominantly affects a set of 5’-TOP motif-containing mRNAs, including those encoding RPs and other components of the translational machinery in cancer cells. However, the detailed molecular mechanism underlying this regulation requires further investigation.

**Figure 4:**
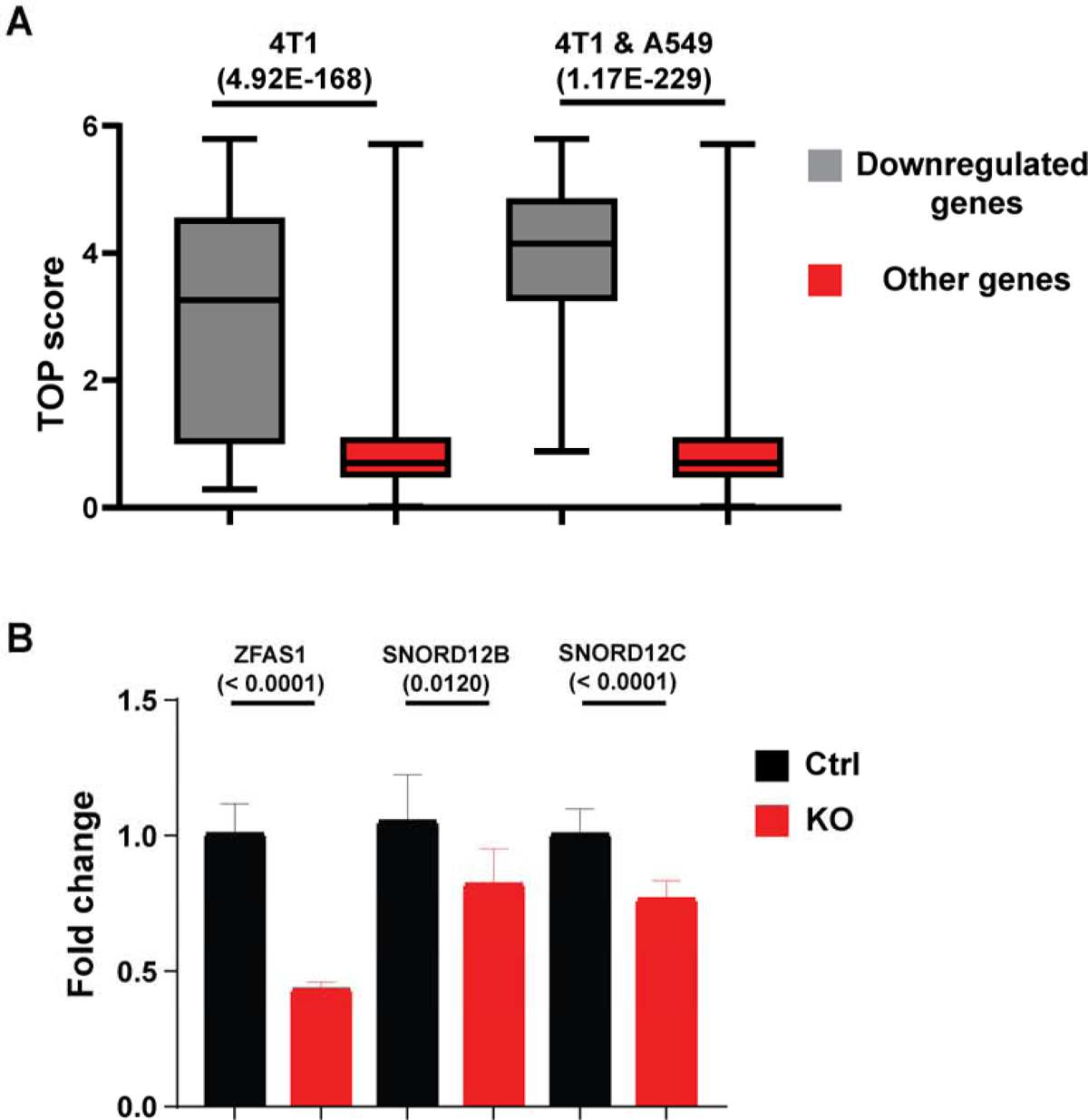
CMTR1 inhibition affects 5’ TOP mRNAs and snoRNA expression in cancer cells. (A) 5’ TOP scores for downregulated genes in 4T1 CMTR1-KO cells (4T1)or the common gene set as shown in Figure 3B compared to all other genes. (B) Relative expression levels of the snoRNA host gene ZFAS1 and its associated snoRNAs (SNORD12B and SNORD12C) measured by qRT-PCR in 4T1 cells after CMTR1 knockout. *p*-values for each RNA, determined by Welch’s t-test, are displayed. Ctrl: Control; KO: CMTR1 knockout. Error bars: SD.

### Downregulation of snoRNA host genes and snoRNAs following CMTR1 depletion

SnoRNAs and their host genes play important roles in several biological processes, including rRNA modification and ribosome biosynthesis [38]. In the human genome, most snoRNA genes are located within the introns of other ‘host’ genes, which can be either protein-coding or non-coding. Interestingly, our integrated RNA-seq analysis revealed that CMTR1 depletion in cancer cells led to the downregulation of five snoRNA non-coding host genes (*SNHG1*, *SNHG8*, *SNHG12*, *GAS5*, *ZFAS1*) and 21 snoRNA protein-coding host genes (e.g., RPS11, RPL39) (Supplementary Table 6A). To validate these findings, we performed qRT-PCR assays to measure the expression of snoRNA non-coding host genes in our CMTR1-depleted breast cancer cells. Our results confirmed that CMTR1 depletion induced the downregulation of these host genes, including *SNHG8*, *ZFAS1*, and *GAS5* (Figure 3C, 4B). Of particular interest, ZFAS1, which encodes three C/D box SNORD12 family members (SNORD12, SNORD12B, and SNORD12C), is significantly overexpressed in various human cancers [39–41]. Our qRT-PCR assays further showed that two snoRNAs, SNORD12B and SNORD12C, were significantly downregulated in our CMTR1-KO 4T1 cells compared to controls (Figure 4B). These data suggest that CMTR1 overexpression in cancer cells likely supports cell growth and cancer phenotypes through multiple mechanisms, including the regulation of both protein-coding and non-coding gene expression.

### *In silico* and *in vitro* screening for novel CMTR1 inhibitors

To further investigate the potential of CMTR1 as a therapeutic target in cancer, we sought to identify novel inhibitors of this enzyme using a combination of in silico and *in vitro* approaches. We used UCSF-Chimera and Autodock software to optimize the 1.9-Å crystal structure of CMTR1 enzymatic domain (PDB: 4N49) for virtual screening (Figure 5A) [42–45]. Three NCI compound sets were screened: the Diversity set (derived from ∼140,000 compounds), Mechanistic set (derived from 37,836 compounds), and Natural Products set (selected from ∼140,000 compounds) [42]. From this initial screen, 18 compounds were selected for validation using Bio-Layer Interferometry (BLI) assays. Our analysis identified three lead compounds— N97911, N169774, and N627666—that demonstrated potential to block CMTR1-m7G RNA binding (Figure 5B). Further examination of their binding modes to the CMTR1 enzymatic domain revealed that all three compounds exhibit hydrophobic interactions and/or π-π interactions with key residue(s) of the m7G RNA pocket (Figure 5C). Notably, data from the NCI Developmental Therapeutics Program indicated that N97911 displayed substantial growth inhibition activity against a set of NCI-60 cancer lines, including ovarian cancer lines (data not shown). We subsequently confirmed that these CMTR1 candidate compounds, particularly N97911, inhibited 4T1 cancer cell growth and survival *in vitro* (Figure 5D). While further studies are necessary, our identified CMTR1 inhibitor candidates provide starting points to identify more potent inhibitors and suggest the therapeutic potential of targeting CMTR1 in various cancers.

**Figure 5.**
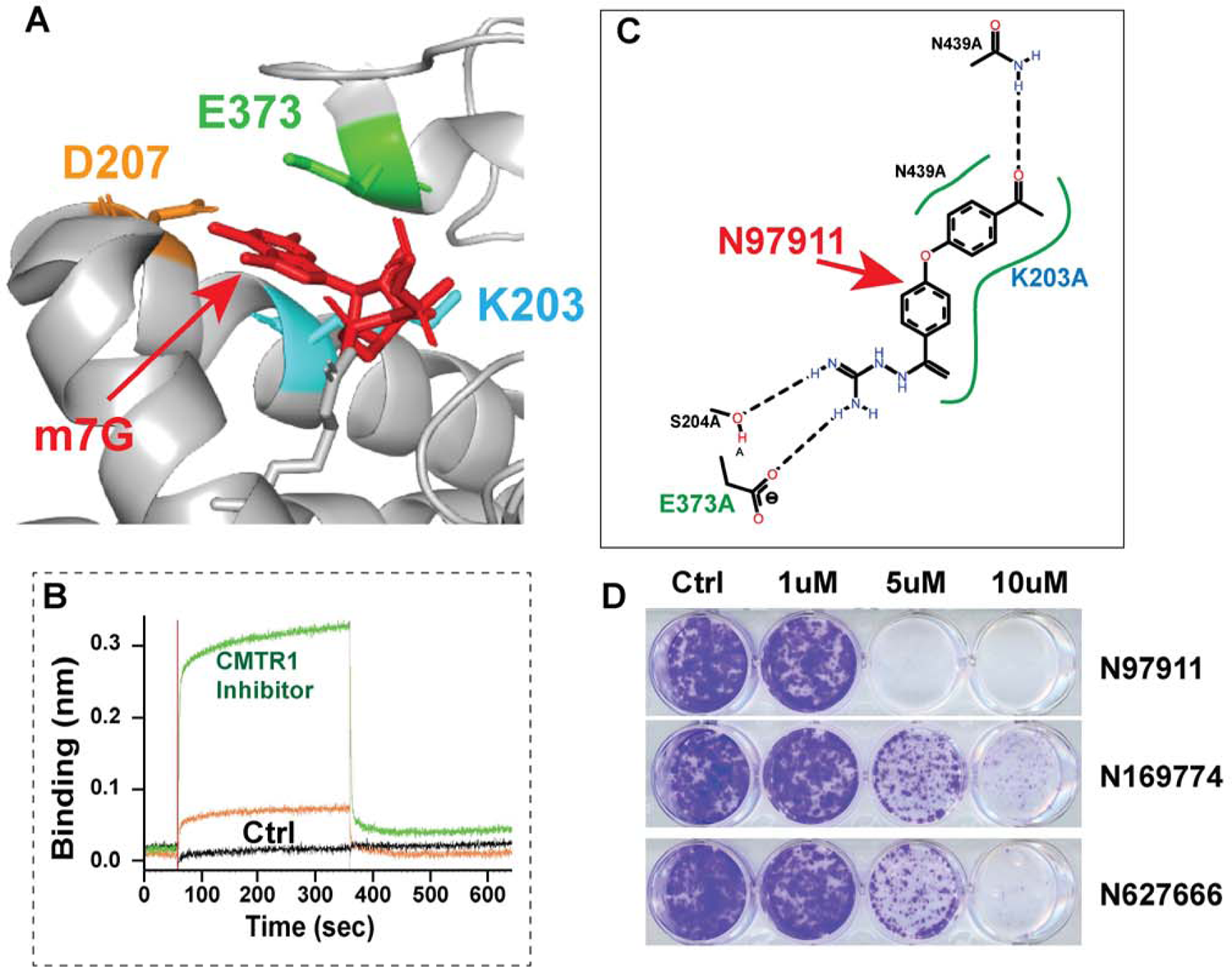
Identification of novel CMTR1 inhibitors. (A) Ribbon diagram showing key residues of CMTR1-Ftsj domain (PDB: 4N49) interacting with m7G RNA. (B) Association/dissociation binding curves (BLI assay) of CMTR1 candidate inhibitors to the immobilized CMTR1-Ftsj protein. (C) 2D view of the predicted N97911-CMTR1 binding mode created by Pose View. (D) Representative images of clonogenic survival in 4T1 cells after treatment with CMTR1 candidate inhibitors.

## Discussion

In this study, we employed a multi-omics approach, combining transcriptomics, proteomics, and functional genomics to elucidate the clinical and biological significance of cap methyltransferases, focusing primarily on CMTR1, in various human cancers. Our analysis revealed differential expression of CMTR1 and CMTR2 across cancer types, with CMTR1, but not CMTR2, showing upregulation at both mRNA and protein levels in several cancer types examined. Functionally, CMTR1 depletion significantly inhibited tumor growth both *in vitro* and *in vivo*. At the molecular level, we found that CMTR1 regulates the expression of ribosomal protein genes and other transcripts with 5’ TOP motifs. Additionally, CMTR1 affects the expression of snoRNA host genes and snoRNAs, suggesting a broader role in RNA metabolism. Finally, we explored the potential of CMTR1 as a therapeutic target by identifying candidate inhibitors. While further studies are necessary, these CMTR1 inhibitor candidates provide promising starting points for developing more potent inhibitors and highlight the therapeutic potential of targeting CMTR1 in various cancers.

CMTR1 was originally identified in 2008 as an interferon-stimulated gene (ISG), initially designated as ISG95, whose expression increases in response to interferon treatment and viral infection [16]. This early study also revealed that CMTR1 interacts with the C-terminal domain (CTD) of RNA polymerase II, suggesting CMTR1’s potential role in regulating gene expression and its involvement in mRNA processing events. Subsequently, Bélanger *et al.* biochemically characterized CMTR1 and revealed that it functions as the 2’-O-ribose methyltransferase responsible for cap1 formation in higher eukaryotic cells [14]. In contrast, its homolog CMTR2, methylates the 2’-O-ribose of the second transcribed nucleotide, forming cap2 structures [13]. Structurally, in addition to the Ftsj methyltransferase domain, CMTR1 contains multiple non-enzymatic domains or motifs, including an N-terminal nuclear localization sequence, a G-patch domain, a C-terminal inactive cap guanylyltransferase-like (GTase-like) domain, and a WW domain [4, 14–16, 19]. A very recent study identified that CMTR1 also contains a highly phosphorylated ‘P-Patch’ motif, which is targeted by the kinase CK2 (casein kinase II) [19]. In contrast, CMTR2 contains only the enzymatic Ftsj methyltransferase domain and a catalytically inactive methyltransferase domain. [12].

Previous studies revealed that the non-enzymatic domains of CMTR1 interact with various proteins, influencing both the enzymatic activity of CMTR1 and transcriptional regulation [14–16]. Early studies revealed that the WW domain of CMTR1 interacts with the Ser-5 phosphorylated C-terminal domain of RNA Pol II [15, 16]. Two recent structural studies of human co-transcriptional capping demonstrated that the C-terminal GTase-like domain of CMTR1 also directly interacts with Pol II, likely playing a role in the recruitment of CMTR1 to the Pol II surface [46, 47]. Furthermore, these studies revealed that CMTR1 directly binds to the paused elongation complex, which includes DRB sensitivity-inducing factor (DSIF) and negative elongation factor (NELF) [46, 47]. These structural studies revealed the co-existence of CMTR1 with pausing factors, suggesting an intricate relationship between 5’ end capping modifications and the transcriptional pausing of Pol II.

CMTR1 also contains a G-patch, a glycine-rich 50-residue motif found in approximately 20 human proteins [15]. Notably, CMTR1 is the only human G-patch protein that possesses a catalytic domain [48]. Among these G-patch proteins, more than ten, including CMTR1, have been identified to interact with the OB-fold (oligonucleotide/oligosaccharide-binding fold) of the RNA helicase DHX15 (DEAH-Box Helicase 15), with most of these interactions enhancing DHX15’s enzymatic activity [48]. Regarding the CMTR1-DHX15 interaction, Inesta-Vaquera et al. suggest that CMTR1-DHX15 interactions reduce the methyltransferase activity of CMTR1 while increasing the helicase activity of DHX15 [15]. In contrast, Toczydlowska-Socha et al. propose that DHX15 facilitates CMTR1’s methyltransferase activity on mRNAs with highly structured 5’ ends, while CMTR1 enhances DHX15’s ATPase activity, though not its helicase activity. DHX15 is a multifunctional RNA helicase involved in RNA splicing, ribosome biogenesis, viral RNA sensing, and other cellular processes [49]. Analysis of CPTAC and CCLE cancer datasets revealed that the DHX15 protein is highly expressed in cancer samples compared to normal tissues and is significantly positively correlated with CMTR1 protein expression (Supplementary Figure 9). These protein correlation data provide additional evidence of the positive interconnection and interplay between CMTR1 and DHX15. However, further mechanistic studies are needed to fully understand how these proteins work together to promote cancer progression.

Recent studies using knockout mice for CMTR1 and CMTR2 revealed that the loss of either CMTR1 or CMTR2 results in lethality; however, the sets of misregulated transcripts do not overlap [50, 51]. Using CMTR1-conditional knockout mice, Lee *et al.* found that CMTR1-catalyzed 2’-O-ribose methylation controls neuronal development by regulating Camk2α (Calcium/Calmodulin Dependent Protein Kinase II Alpha) expression [52]. Studies also demonstrated that CMTR1 plays an important role in the differentiation of embryonic stem cells by promoting the expression of ribosomal proteins and histone genes [17, 53]. Notably, our RNA-seq analysis in various cancer models also revealed that CMTR1 regulates the expression of ribosomal protein genes but not histone genes. This suggests that ribosomal protein genes are commonly regulated downstream of CMTR1, possibly through CMTR1 recognition of transcripts with 5’ TOP motifs.

Prior to our study, research on CMTR1 in cancer was limited but provided initial insights into its potential roles. In 2018, Du *et al.* reported a CMTR1-ALK fusion in non-small-cell lung cancer that did not respond to the ALK inhibitor crizotinib, suggesting a possible role for CMTR1 in drug resistance [21]. More recently, You *et al.* demonstrated that CMTR1 promotes colorectal cancer cell growth and immune evasion by transcriptionally regulating STAT3 (signal transducer and activator of transcription 3) [22]. They showed that CMTR1 knockdown reduced colon cancer cell proliferation and tumor growth *in vivo*, while also enhancing the efficacy of anti-PD-1 therapy [22]. Previous studies have also revealed that CMTR1 knockdown inhibits *in vitro* cell proliferation of human MCF7 and HCC1806 breast cancer cell lines [15]. Our work advances our understanding of CMTR1 in cancer, as well as a set of common genes (especially ribosomal) regulated by CMTR1 across tumor types.

In conclusion, our comprehensive study establishes CMTR1 as a critical player in cancer biology, demonstrating its widespread upregulation across multiple tumor types and significant impacts on cell growth, ribosomal protein gene expression, and snoRNA regulation. Our discovery of a potential CMTR1 inhibitor opens promising avenues for therapeutic intervention. While these findings advance our understanding of CMTR1 in cancer, they also highlight the need for further research to fully elucidate the protein-level effects of CMTR1 depletion, its impact on ribosomal function, and its interplay with other cap methyltransferases. As we continue to unravel the complexities of CMTR1’s role in cancer, this enzyme emerges as a promising target for novel therapies in cancer and other diseases, such as viral infections.

## Materials and Methods

### Bioinformatic data collection and analyses

Normalized RNA-sequencing data from 11,069 TCGA samples, including 737 normal samples, were downloaded from the GDC portal (https://gdc.cancer.gov). Normalized proteomics, phosphoproteomics, and/or acetylproteome data from CPTAC samples were downloaded from LinkedOmics (http://linkedomics.org) [54, 55]. Tumor types containing at least 10 paired TCGA or CPTAC normal samples were selected to calculate the mRNA and protein expression differences between tumor and normal samples. Clinical information was also downloaded from LinkedOmics. Statistical differences in gene, protein, or modified protein levels were calculated by Mann-Whitney U test or Brown-Forsythe and Welch ANOVA using GraphPad Prism 10 or R. Phosphorylation/protein ratio data were queried using CProSite (https://cprosite.ccr.cancer.gov/) [28]. Cancer dependency scores for CMTR1 and CMTR2 were downloaded from the DepMap website (https://depmap.org/portal/) using the 24Q2 dataset [33]. For RNA-seq data (GSE141171) analysis of A549 CMTR1 WT and KO cells with or without IFN treatment, differential gene expression was performed using GREIN (http://www.ilincs.org/apps/grein/?gse<u>=</u>) [56].

### CRISPR-mediated CMTR1 knockout and siRNA CMTR1 knockdown

The CRISPR-mediated CMTR1 knockout 4T1 model was generated by Synthego. Briefly, we designed a single guide RNA (sgRNA) with the target sequence AUUCGCUUCUGUUUCUUGAG, to knock out mouse CMTR1. This sgRNA was complexed with SpCas9 (Streptococcus pyogenes Cas9) to create a ribonucleoprotein (RNP). The RNP was then introduced into the 4T1 cells using optimized electroporation settings. Subsequently, we employed PCR and Sanger sequencing to confirm the successful knockout of CMTR1 in the 4T1 cells. The CMTR1 protein levels in 4T1 CRISPR control and CMTR1-KO cells were also measured by Western blotting. For siRNA knockdown of CMTR1 in the human MDA-MB-436 breast cancer model, cells were plated in 6-well plates (for RNA and protein collection) or 24-well plates (for survival assay) at proportional concentrations. Cells were then transfected using Sigma-Aldrich MISSION esiRNAs targeting CMTR1 or control eGFP according to the manufacturer’s protocol. RNA was collected 48 hours after transfection, while protein was collected 72 or 96 hours post-transfection. Cell proliferation and/or survival in CRISPR-mediated CMTR1 knockout 4T1 cells and siRNA CMTR1 knockdown MDA-MB-436 cells were measured using a CellTiter-Blue Cell Viability Assay Kit and crystal violet staining.

### Soft agar assay

Soft agar assays were performed as previously described [57]. Briefly, dishes were coated with a 1:1 mixture of the appropriate 2× medium for the 4T1 cell line and 1% Bacto agar. Cells were plated at 1 × 10^4^ per well, fed three times per week for 3 to 4 weeks, stained overnight with 500 μg/mL p-iodonitrotetrazolium violet (Sigma, St. Louis, MO), and counted using an Oxford Optronix GelCount colony counter.

### Analysis of 4T1 RNA-Seq Data

For RNA-seq of 4T1 WT and CMTR1-KO cells, poly(A) RNA sequencing libraries were prepared following Illumina’s TruSeq-stranded-mRNA sample preparation protocol. RNA integrity was checked with an Agilent Technologies 2100 Bioanalyzer. Poly(A) tail-containing mRNAs were purified using oligo-(dT) magnetic beads with two rounds of purification. After purification, poly(A) RNA was fragmented using divalent cation buffer at elevated temperature. Quality control analysis and quantification of the sequencing libraries were performed using an Agilent Technologies 2100 Bioanalyzer High Sensitivity DNA Chip. Paired-end sequencing was performed on Illumina’s NovaSeq 6000 sequencing system. We used HISAT2 to map reads to the mouse genome (mm10)). Differentially expressed mRNAs were identified using the R package edgeR. Gene Ontology analysis was performed with Enrichr using the KEGG 2019 Mouse or KEGG 2021 Human datasets (https://maayanlab.cloud/Enrichr/) [58].

### RNA preparation and semiquantitative PCR reactions

To assess gene expression changes at the mRNA level, RNA was extracted from 4T1 and MDA-MB-436 cell lines using a RNeasy Plus Mini Kit (QIAGEN). The RNA was mixed with qScript cDNA SuperMix (Quanta Biosciences, Gaithersburg, MD, USA) and converted to cDNA through reverse transcription (RT). The resulting cDNA was then used for real-time PCR reactions with the following settings: 50°C for 2 minutes, 95°C for 10 minutes, 40 or 45 cycles of 95°C for 15 seconds, 60°C for 30 seconds, 72°C for 30 seconds, followed by 72°C for 10 minutes. Sequences of a set of primers for genes and snoRNAs were obtained from PrimerBank, or designed using Primer3 tool [59, 60]. PUM1 (human) or GAPDH (mouse) primer sets were used as controls.

### *In vivo* tumor growth

All animal studies were approved by the Wayne State University Institutional Animal Care and Use Committee. A total of 2 × 10 wild-type (WT) or CMTR1-KO 4T1 cells were injected into the mammary fat pads of female BALB/c mice. Tumor size was measured with calipers two to three times per week, and mice were euthanized when the tumor burden exceeded 1,500 mg. Bilateral tumor volumes from individual mice were used to calculate the growth of WT and CMTR1-KO tumors using the formula volume (mg) = L x w^2^/2 where length (L, mm) and width (w) were determined by caliper measurements.

### Virtual and biochemical screening of CMTR1 inhibitor

The crystal structure of human CMTR1 (PDB: 4N49) at 1.9 Å resolution was prepared with UCSF Chimera for virtual screening against three NCI compound sets: the Diversity Set, Mechanistic Set, and Natural Products Set [61, 62]. For MTiOpenScreen Vina docking, the grid center coordinates for the SAM binding pocket of 4N49 were set to (x, y, z) = (8.9, 20.8, 16.6), with a search space size of 20 Å x 20 Å x 20 Å. The MTiOpenScreen screening was repeated three times, and compounds shared across all three runs were further analyzed using DataWarrior, PyMOL, and UCSF Chimera. Based on virtual screening scores and predicted physicochemical properties, 18 compounds were selected for future validation using Bio-Layer Interferometry (BLI) assays, which were performed according to the manufacturer’s protocol.

## Acknowledgments

This work was partially supported by grants from the Department of Defense (DoD) Breast Cancer Program BC201476, Elsa U. Pardee Foundation, DMC Foundation and Molecular Therapeutics Program of Karmanos Cancer Institute to Dr. Zeng-Quan Yang. The Animal Model and Therapeutics Evaluation, and Proteomics Cores were supported in part by NIH Center grant P30 CA22453 to the Barbara Ann Karmanos Cancer Institute. We thank Paul Stemmer, Lanxin Liu, Rui Wang, Juiwanna Kushner, Maksymilian Pilecki, Ivan Lopez and Morenci Manning for technical contributions.

## Supplementary Figure Legends

**Supplementary Figure 1:** mRNA and protein expression levels of CMTR2 in tumor and normal adjacent tissue (NAT) samples across TCGA and CPTAC datasets. (A) Dot plots of log2(RSEM) mRNA expression values of CMTR2 in TCGA dataset. (B) Dot plots of log2 TMT protein abundance of CMTR2 in CPTAC dataset. Blue dots represent NAT samples, whereas red dots represent tumor samples. Each data point represents an individual patient sample. The p-value from Mann-Whitney U test comparing tumor and NAT samples for each cancer type is displayed. Error bars indicate standard deviation of mRNA or protein expression levels across patients. RSEM: RNA-Seq by Expectation-Maximization; TMT: Tandem Mass Tags; TCGA: The Cancer Genome Atlas; CPTAC: Clinical Proteomic Tumor Analysis Consortium.

**Supplementary Figure 2:** Expression differences (log2 FC) in CMTR1 protein phosphorylation and acetylation levels between CPTAC tumor and normal adjacent tissue (NAT) samples. The heatmap displays log2 FC for cancer types with at least 10 samples in both tumor and NAT groups. The color gradient from purple to red represents the degree of log2 FC between tumor and normal samples. Dot size indicates statistical significance. Only data points with log FC >|0.2| and p-value < 0.05 are shown. FC: fold-change.

**Supplementary Figure 3:** CMTR1 shows increased expression in serous and CNV-high subtypes and correlates with reduced overall survival (OS) in UCEC CPTAC and TCGA data. (A) Protein abundance of CMTR1 in NAT, endometrioid, and serous UCEC samples from CPTAC proteomic data. (B) Kaplan-Meier overall survival (OS) curves of TCGA UCEC patients (n = 513) stratified into quartiles by CMTR1 expression. The highest expression group (Q4) is shown with a red line. (C) Protein abundance of CMTR1 in NAT and different molecular subtypes (CNV-high, CNV-low, MSI, and POLE) of UCEC from CPTAC proteomic data. CNV: copy number variation; MSI, microsatellite instability; POLE, DNA polymerase epsilon.

**Supplementary Figure 4:** CMTR1 and CMTR2 dependency scores in cancer cell lines and CMTR1 expression across breast cancer subtypes. (A) Dot plots showing dependency scores of CMTR1 and CMTR2 in genome-scale loss-of-function screens (DepMap 24Q2) across 1,150 cancer cell lines. (B) mRNA expression levels of CMTR1 across five subtypes of TCGA breast cancer samples: normal-like, luminal A, luminal B, HER2-enriched, and basal-like. DepMap: Cancer Dependency Map; HER2, human epidermal growth factor receptor 2.

**Supplementary Figure 5:** Relative higher expression of CMTR1 protein in a subset of basal-like breast cancer cell lines. (A) Relative protein abundance of CMTR1 in 30 breast cancer cell lines from the Cancer Cell Line Encyclopedia (CCLE). The cell lines are categorized into five subtypes: normal-like, luminal A, luminal B, HER2-enriched, and basal-like. (B) CMTR1 protein levels were assessed by Western blot assay in a panel of breast cancer cell lines. The cell lines are color-coded as follows: normal-like (green), Luminal A: light blue, luminal B: dark blue, HER2-enriched (purple), and basal-like (red).

**Supplementary Figure 6:** Inhibition of CMTR1 suppresses 4T1 breast cancer cell invasion. The effect of CMTR1 knockout (KO) on the invasion potential of 4T1 cells was assessed using a Transwell assay. Cells were stained with calcein-AM, and images of the stained invaded cells were captured under a fluorescent microscope. Representative images at 4x magnification of wild-type (WT, top) and CMTR1 KO (bottom) cells from invasion assays are shown.,

**Supplementary Figure 7:** CMTR1 knockout reduces expression of ribosomal protein genes and SNHGs in A549 cells. Volcano plot of up-and down-regulated genes after CMTR1 knockout in A549 NT (top) and A549 IFN-treated (bottom) cells. Significanltly up- or down-regulated genes with q < 0.05 is indicated in green, ribosomal or SNHG gene is indicated in red. Significance is based on q-value < 0.05, as determined by Benjamini-Hochberg correction of calculated *p*-values.

**Supplementary Figure 8:** CMTR1 knockdown reduces expression of ribosomal protein and small nucleolar RNA host genes in basal-like breast cancer MDA-MB-231 cells. Bar graphs show qRT-PCR results of indicated mRNAs from MDA-MB-231 cells with or without CMTR1 knockdown. Data were normalized to PUM1 and are presented as fold change relative to the control. p-values for each gene, determined by Welch’s t-test, are displayed. Error bars: SD; SNHG: small nucleolar RNA host gene.

**Supplementary Figure 9:** DHX15 protein abundance is significantly positively correlated with CMTR1 protein abundance in 1,301 CPTAC tumor samples.

## References

[1] R.G. Avila-Bonilla, S. Macias, The molecular language of RNA 5’ ends: guardians of RNA identity and immunity, RNA, 30 (2024) 327–336.

[2] A. Ramanathan, G.B. Robb, S.H. Chan, mRNA capping: biological functions and applications, Nucleic Acids Res, 44 (2016) 7511–7526.

[3] F. Inesta-Vaquera, V.H. Cowling, Regulation and function of CMTR1-dependent mRNA cap methylation, Wiley Interdiscip Rev RNA, 8 (2017).

[4] S. Liang, R. Almohammed, V.H. Cowling, The RNA cap methyltransferases RNMT and CMTR1 co-ordinate gene expression during neural differentiation, Biochem Soc Trans, 51 (2023) 1131–1141.

[5] K. Borden, B. Culjkovic-Kraljacic, V.H. Cowling, To cap it all off, again: dynamic capping and recapping of coding and non-coding RNAs to control transcript fate and biological activity, Cell Cycle, 20 (2021) 1347–1360.

[6] J. Pelletier, T.M. Schmeing, N. Sonenberg, The multifaceted eukaryotic cap structure, Wiley Interdiscip Rev RNA, 12 (2021) e1636.

[7] V.H. Cowling, Regulation of mRNA cap methylation, Biochem J, 425 (2009) 295–302.

[8] T. Yamada-Okabe, R. Doi, O. Shimmi, M. Arisawa, H. Yamada-Okabe, Isolation and characterization of a human cDNA for mRNA 5’-capping enzyme, Nucleic Acids Res, 26 (1998) 1700–1706.

[9] H. Bugl, E.B. Fauman, B.L. Staker, F. Zheng, S.R. Kushner, M.A. Saper, J.C. Bardwell, U. Jakob, RNA methylation under heat shock control, Mol Cell, 6 (2000) 349–360.

[10] K.I. Zhou, C.V. Pecot, C.L. Holley, 2’-O-methylation (Nm) in RNA: progress, challenges, and future directions, RNA, 30 (2024) 570–582.

[11] P.O. Falnes, Closing in on human methylation-the versatile family of seven-beta-strand (METTL) methyltransferases, Nucleic Acids Res, (2024).

[12] M. Smietanski, M. Werner, E. Purta, K.H. Kaminska, J. Stepinski, E. Darzynkiewicz, M. Nowotny, J.M. Bujnicki, Structural analysis of human 2’-O-ribose methyltransferases involved in mRNA cap structure formation, Nat Commun, 5 (2014) 3004.

[13] M. Werner, E. Purta, K.H. Kaminska, I.A. Cymerman, D.A. Campbell, B. Mittra, J.R. Zamudio, N.R. Sturm, J. Jaworski, J.M. Bujnicki, 2’-O-ribose methylation of cap2 in human: function and evolution in a horizontally mobile family, Nucleic Acids Res, 39 (2011) 4756–4768.

[14] F. Belanger, J. Stepinski, E. Darzynkiewicz, J. Pelletier, Characterization of hMTr1, a human Cap1 2’-O-ribose methyltransferase, J Biol Chem, 285 (2010) 33037–33044.

[15] F. Inesta-Vaquera, V.K. Chaugule, A. Galloway, L. Chandler, A. Rojas-Fernandez, S. Weidlich, M. Peggie, V.H. Cowling, DHX15 regulates CMTR1-dependent gene expression and cell proliferation, Life Sci Alliance, 1 (2018) e201800092.

[16] T. Haline-Vaz, T.C. Silva, N.I. Zanchin, The human interferon-regulated ISG95 protein interacts with RNA polymerase II and shows methyltransferase activity, Biochem Biophys Res Commun, 372 (2008) 719–724.

[17] S. Liang, J.C. Silva, O. Suska, R. Lukoszek, R. Almohammed, V.H. Cowling, CMTR1 is recruited to transcription start sites and promotes ribosomal protein and histone gene expression in embryonic stem cells, Nucleic Acids Res, 50 (2022) 2905–2922.

[18] B. Li, S.M. Clohisey, B.S. Chia, B. Wang, A. Cui, T. Eisenhaure, L.D. Schweitzer, P. Hoover, N.J. Parkinson, A. Nachshon, N. Smith, T. Regan, D. Farr, M.U. Gutmann, S.I. Bukhari, A. Law, M. Sangesland, I. Gat-Viks, P. Digard, S. Vasudevan, D. Lingwood, D.H. Dockrell, J.G. Doench, J.K. Baillie, N. Hacohen, Genome-wide CRISPR screen identifies host dependency factors for influenza A virus infection, Nat Commun, 11 (2020) 164.

[19] R. Lukoszek, F. Inesta-Vaquera, N.J.M. Brett, S. Liang, L.A. Hepburn, D.J. Hughes, C. Pirillo, E.W. Roberts, V.H. Cowling, CK2 phosphorylation of CMTR1 promotes RNA cap formation and influenza virus infection, Cell Rep, 43 (2024) 114405.

[20] G.D. Williams, N.S. Gokhale, D.L. Snider, S.M. Horner, The mRNA Cap 2’-O-Methyltransferase CMTR1 Regulates the Expression of Certain Interferon-Stimulated Genes, mSphere, 5 (2020).

[21] X. Du, Y. Shao, H. Gao, X. Zhang, H. Zhang, Y. Ban, H. Qin, Y. Tai, CMTR1-ALK: an ALK fusion in a patient with no response to ALK inhibitor crizotinib, Cancer Biol Ther, 19 (2018) 962–966.

[22] A.B. You, H. Yang, C.P. Lai, W. Lei, L. Yang, J.L. Lin, S.C. Liu, N. Ding, F. Ye, CMTR1 promotes colorectal cancer cell growth and immune evasion by transcriptionally regulating STAT3, Cell Death Dis, 14 (2023) 245.

[23] D.G. Dimitrova, L. Teysset, C. Carre, RNA 2’-O-Methylation (Nm) Modification in Human Diseases, Genes (Basel), 10 (2019).

[24] M. Manning, Y. Jiang, R. Wang, L. Liu, S. Rode, M. Bonahoom, S. Kim, Z.Q. Yang, Pan-cancer analysis of RNA methyltransferases identifies FTSJ3 as a potential regulator of breast cancer progression, Rna Biol, 17 (2020) 474–486.

[25] Y. Li, Y. Dou, F. Da Veiga Leprevost, Y. Geffen, A.P. Calinawan, F. Aguet, Y. Akiyama, S. Anand, C. Birger, S. Cao, R. Chaudhary, P. Chilappagari, M. Cieslik, A. Colaprico, D.C. Zhou, C. Day, M.J. Domagalski, M. Esai Selvan, D. Fenyo, S.M. Foltz, A. Francis, T. Gonzalez-Robles, Z.H. Gumus, D. Heiman, M. Holck, R. Hong, Y. Hu, E.J. Jaehnig, J. Ji, W. Jiang, L. Katsnelson, K.A. Ketchum, R.J. Klein, J.T. Lei, W.W. Liang, Y. Liao, C.M. Lindgren, W. Ma, L. Ma, M.J. MacCoss, F. Martins Rodrigues, W. McKerrow, N. Nguyen, R. Oldroyd, A. Pilozzi, P. Pugliese, B. Reva, P. Rudnick, K.V. Ruggles, D. Rykunov, S.R. Savage, M. Schnaubelt, T. Schraink, Z. Shi, D. Singhal, X. Song, E. Storrs, N.V. Terekhanova, R.R. Thangudu, M. Thiagarajan, L.B. Wang, J.M. Wang, Y. Wang, B. Wen, Y. Wu, M.A. Wyczalkowski, Y. Xin, L. Yao, X. Yi, H. Zhang, Q. Zhang, M. Zuhl, G. Getz, L. Ding, A.I. Nesvizhskii, P. Wang, A.I. Robles, B. Zhang, S.H. Payne, C. Clinical Proteomic Tumor Analysis, Proteogenomic data and resources for pan-cancer analysis, Cancer Cell, 41 (2023) 1397–1406.

[26] C.T. Walsh, S. Garneau-Tsodikova, G.J. Gatto, Jr., Protein posttranslational modifications: the chemistry of proteome diversifications, Angew Chem Int Ed Engl, 44 (2005) 7342–7372.

[27] P.V. Hornbeck, J.M. Kornhauser, V. Latham, B. Murray, V. Nandhikonda, A. Nord, E. Skrzypek, T. Wheeler, B. Zhang, F. Gnad, 15 years of PhosphoSitePlus(R): integrating post-translationally modified sites, disease variants and isoforms, Nucleic Acids Res, 47 (2019) D433–D441.

[28] D. Wang, X. Qian, Y.N. Du, B. Sanchez-Solana, K. Chen, M. Kanigicherla, L.M. Jenkins, J. Luo, S. Eng, B. Park, B. Chen, X. Yao, T. Nguyen, B.K. Tripathi, M.E. Durkin, D.R. Lowy, cProSite: A web based interactive platform for online proteomics, phosphoproteomics, and genomics data analysis, J Biotechnol Biomed, 6 (2023) 573–578.

[29] M.E. Urick, D.W. Bell, Clinical actionability of molecular targets in endometrial cancer, Nat Rev Cancer, 19 (2019) 510–521.

[30] V.W. Setiawan, H.P. Yang, M.C. Pike, S.E. McCann, H. Yu, Y.B. Xiang, A. Wolk, N. Wentzensen, N.S. Weiss, P.M. Webb, P.A. van den Brandt, K. van de Vijver, P.J. Thompson, G. Australian National Endometrial Cancer Study, B.L. Strom, A.B. Spurdle, R.A. Soslow, X.O. Shu, C. Schairer, C. Sacerdote, T.E. Rohan, K. Robien, H.A. Risch, F. Ricceri, T.R. Rebbeck, R. Rastogi, J. Prescott, S. Polidoro, Y. Park, S.H. Olson, K.B. Moysich, A.B. Miller, M.L. McCullough, R.K. Matsuno, A.M. Magliocco, G. Lurie, L. Lu, J. Lissowska, X. Liang, J.V. Lacey, Jr., L.N. Kolonel, B.E. Henderson, S.E. Hankinson, N. Hakansson, M.T. Goodman, M.M. Gaudet, M. Garcia-Closas, C.M. Friedenreich, J.L. Freudenheim, J. Doherty, I. De Vivo, K.S. Courneya, L.S. Cook, C. Chen, J.R. Cerhan, H. Cai, L.A. Brinton, L. Bernstein, K.E. Anderson, H. Anton-Culver, L.J. Schouten, P.L. Horn-Ross, Type I and II endometrial cancers: have they different risk factors?, J Clin Oncol, 31 (2013) 2607–2618.

[31] N. Cancer Genome Atlas Research, C. Kandoth, N. Schultz, A.D. Cherniack, R. Akbani, Y. Liu, H. Shen, A.G. Robertson, I. Pashtan, R. Shen, C.C. Benz, C. Yau, P.W. Laird, L. Ding, W. Zhang, G.B. Mills, R. Kucherlapati, E.R. Mardis, D.A. Levine, Integrated genomic characterization of endometrial carcinoma, Nature, 497 (2013) 67–73.

[32] G. Bogani, I. Ray-Coquard, N. Concin, N.Y.L. Ngoi, P. Morice, T. Enomoto, K. Takehara, H. Denys, R.A. Nout, D. Lorusso, M.M. Vaughan, M. Bini, M. Takano, D. Provencher, A. Indini, S. Sagae, P. Wimberger, R. Poka, Y. Segev, S.I. Kim, F.J. Candido Dos Reis, S. Lopez, A. Mariani, M.M. Leitao, Jr., F. Raspagliesi, P.B. Panici, V. Di Donato, L. Muzii, N. Colombo, G. Scambia, S. Pignata, B.J. Monk, Uterine serous carcinoma, Gynecol Oncol, 162 (2021) 226–234.

[33] A. Tsherniak, F. Vazquez, P.G. Montgomery, B.A. Weir, G. Kryukov, G.S. Cowley, S. Gill, W.F. Harrington, S. Pantel, J.M. Krill-Burger, R.M. Meyers, L. Ali, A. Goodale, Y. Lee, G. Jiang, J. Hsiao, W.F.J. Gerath, S. Howell, E. Merkel, M. Ghandi, L.A. Garraway, D.E. Root, T.R. Golub, J.S. Boehm, W.C. Hahn, Defining a Cancer Dependency Map, Cell, 170 (2017) 564–576 e516.

[34] T.P. Howard, F. Vazquez, A. Tsherniak, A.L. Hong, M. Rinne, A.J. Aguirre, J.S. Boehm, W.C. Hahn, Functional Genomic Characterization of Cancer Genomes, Cold Spring Harb Symp Quant Biol, 81 (2016) 237–246.

[35] S. Borowicz, M. Van Scoyk, S. Avasarala, M.K. Karuppusamy Rathinam, J. Tauler, R.K. Bikkavilli, R.A. Winn, The soft agar colony formation assay, J Vis Exp, (2014) e51998.

[36] E. Cockman, P. Anderson, P. Ivanov, TOP mRNPs: Molecular Mechanisms and Principles of Regulation, Biomolecules, 10 (2020).

[37] L. Philippe, A.M.G. van den Elzen, M.J. Watson, C.C. Thoreen, Global analysis of LARP1 translation targets reveals tunable and dynamic features of 5’ TOP motifs, Proc Natl Acad Sci U S A, 117 (2020) 5319–5328.

[38] Z.H. Huang, Y.P. Du, J.T. Wen, B.F. Lu, Y. Zhao, snoRNAs: functions and mechanisms in biological processes, and roles in tumor pathophysiology, Cell Death Discov, 8 (2022) 259.

[39] S.J. O’Brien, C. Fiechter, J. Burton, J. Hallion, M. Paas, A. Patel, A. Patel, A. Rochet, K. Scheurlen, S. Gardner, M. Eichenberger, H. Sarojini, S. Srivastava, S. Rai, T. Kalbfleisch, H.C. Polk, Jr., S. Galandiuk, Long non-coding RNA ZFAS1 is a major regulator of epithelial-mesenchymal transition through miR-200/ZEB1/E-cadherin, vimentin signaling in colon adenocarcinoma, Cell Death Discov, 7 (2021) 61.

[40] M. Rao, S. Xu, Y. Zhang, Y. Liu, W. Luan, J. Zhou, Long non-coding RNA ZFAS1 promotes pancreatic cancer proliferation and metastasis by sponging miR-497-5p to regulate HMGA2 expression, Cell Death Dis, 12 (2021) 859.

[41] H. Wu, W. Qin, S. Lu, X. Wang, J. Zhang, T. Sun, X. Hu, Y. Li, Q. Chen, Y. Wang, H. Zhao, H. Piao, R. Zhang, M. Wei, Long noncoding RNA ZFAS1 promoting small nucleolar RNA-mediated 2’-O-methylation via NOP58 recruitment in colorectal cancer, Mol Cancer, 19 (2020) 95.

[42] S. Forli, R. Huey, M.E. Pique, M.F. Sanner, D.S. Goodsell, A.J. Olson, Computational protein-ligand docking and virtual drug screening with the AutoDock suite, Nat Protoc, 11 (2016) 905–919.

[43] X. Fradera, K. Babaoglu, Overview of Methods and Strategies for Conducting Virtual Small Molecule Screening, Curr Protoc Chem Biol, 9 (2017) 196–212.

[44] S. Forli, Charting a Path to Success in Virtual Screening, Molecules, 20 (2015) 18732–18758.

[45] E.F. Pettersen, T.D. Goddard, C.C. Huang, G.S. Couch, D.M. Greenblatt, E.C. Meng, T.E. Ferrin, UCSF Chimera--a visualization system for exploratory research and analysis, J Comput Chem, 25 (2004) 1605–1612.

[46] G. Garg, C. Dienemann, L. Farnung, J. Schwarz, A. Linden, H. Urlaub, P. Cramer, Structural insights into human co-transcriptional capping, Mol Cell, 83 (2023) 2464–2477 e2465.

[47] Y. Li, Q. Wang, Y. Xu, Z. Li, Structures of co-transcriptional RNA capping enzymes on paused transcription complex, Nat Commun, 15 (2024) 4622.

[48] K.E. Bohnsack, R. Ficner, M.T. Bohnsack, S. Jonas, Regulation of DEAH-box RNA helicases by G-patch proteins, Biol Chem, 402 (2021) 561–579.

[49] Q. Feng, K. Krick, J. Chu, C.B. Burge, Splicing quality control mediated by DHX15 and its G-patch activator SUGP1, Cell Rep, 42 (2023) 113223.

[50] M. Dohnalkova, K. Krasnykov, M. Mendel, L. Li, O. Panasenko, F. Fleury-Olela, C.B. Vagbo, D. Homolka, R.S. Pillai, Essential roles of RNA cap-proximal ribose methylation in mammalian embryonic development and fertility, Cell Rep, 42 (2023) 112786.

[51] A.V. Yermalovich, Z. Mohsenin, M. Cowdin, B. Giotti, A. Gupta, A. Feng, L. Golomb, D.B. Wheeler, K. Xu, A. Tsankov, O. Cleaver, M. Meyerson, An essential role for Cmtr2 in mammalian embryonic development, Dev Biol, 516 (2024) 47–58.

[52] Y.L. Lee, F.C. Kung, C.H. Lin, Y.S. Huang, CMTR1-Catalyzed 2’-O-Ribose Methylation Controls Neuronal Development by Regulating Camk2alpha Expression Independent of RIG-I Signaling, Cell Rep, 33 (2020) 108269.

[53] F.M. Simabuco, I.C.B. Pavan, N.F. Pestana, P.C. Carvalho, F.L. Basei, D. Campos Granato, A.F. Paes Leme, N.I.T. Zanchin, Interactome analysis of the human Cap-specific mRNA (nucleoside-2’-O-)-methyltransferase 1 (hMTr1) protein, J Cell Biochem, 120 (2019) 5597–5611.

[54] Y. Liao, S.R. Savage, Y. Dou, Z. Shi, X. Yi, W. Jiang, J.T. Lei, B. Zhang, A proteogenomics data-driven knowledge base of human cancer, Cell Syst, 14 (2023) 777–787 e775.

[55] S.V. Vasaikar, P. Straub, J. Wang, B. Zhang, LinkedOmics: analyzing multi-omics data within and across 32 cancer types, Nucleic Acids Res, 46 (2018) D956–D963.

[56] N.A. Mahi, M.F. Najafabadi, M. Pilarczyk, M. Kouril, M. Medvedovic, GREIN: An Interactive Web Platform for Re-analyzing GEO RNA-seq Data, Sci Rep, 9 (2019) 7580.

[57] Z.Q. Yang, K.L. Streicher, M.E. Ray, J. Abrams, S.P. Ethier, Multiple interacting oncogenes on the 8p11-p12 amplicon in human breast cancer, Cancer Res, 66 (2006) 11632–11643.

[58] Z. Xie, A. Bailey, M.V. Kuleshov, D.J.B. Clarke, J.E. Evangelista, S.L. Jenkins, A. Lachmann, M.L. Wojciechowicz, E. Kropiwnicki, K.M. Jagodnik, M. Jeon, A. Ma’ayan, Gene Set Knowledge Discovery with Enrichr, Curr Protoc, 1 (2021) e90.

[59] X. Wang, A. Spandidos, H. Wang, B. Seed, PrimerBank: a PCR primer database for quantitative gene expression analysis, 2012 update, Nucleic Acids Res, 40 (2012) D1144–1149.

[60] A. Untergasser, I. Cutcutache, T. Koressaar, J. Ye, B.C. Faircloth, M. Remm, S.G. Rozen, Primer3--new capabilities and interfaces, Nucleic Acids Res, 40 (2012) e115.

[61] N. Lagarde, E. Goldwaser, T. Pencheva, D. Jereva, I. Pajeva, J. Rey, P. Tuffery, B.O. Villoutreix, M.A. Miteva, A Free Web-Based Protocol to Assist Structure-Based Virtual Screening Experiments, Int J Mol Sci, 20 (2019).

[62] N. Lagarde, J. Rey, A. Gyulkhandanyan, P. Tuffery, M.A. Miteva, B.O. Villoutreix, Online structure-based screening of purchasable approved drugs and natural compounds: retrospective examples of drug repositioning on cancer targets, Oncotarget, 9 (2018) 32346–32361.

