## Supplemental Figures for "Multi-omics analysis reveals CMTR1 upregulation in cancer and roles in ribosomal protein gene expression and tumor growth"

### Supplementary Figure 1

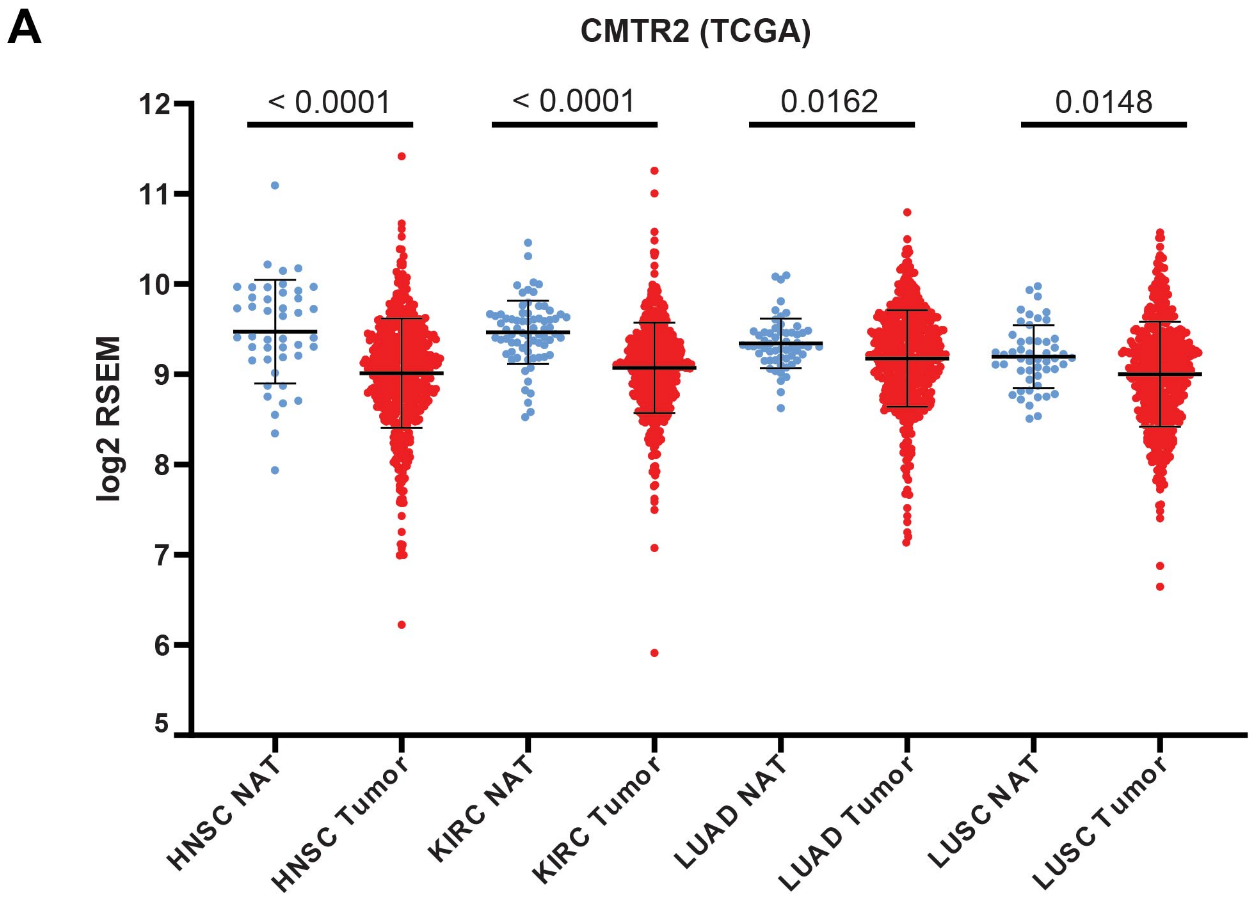


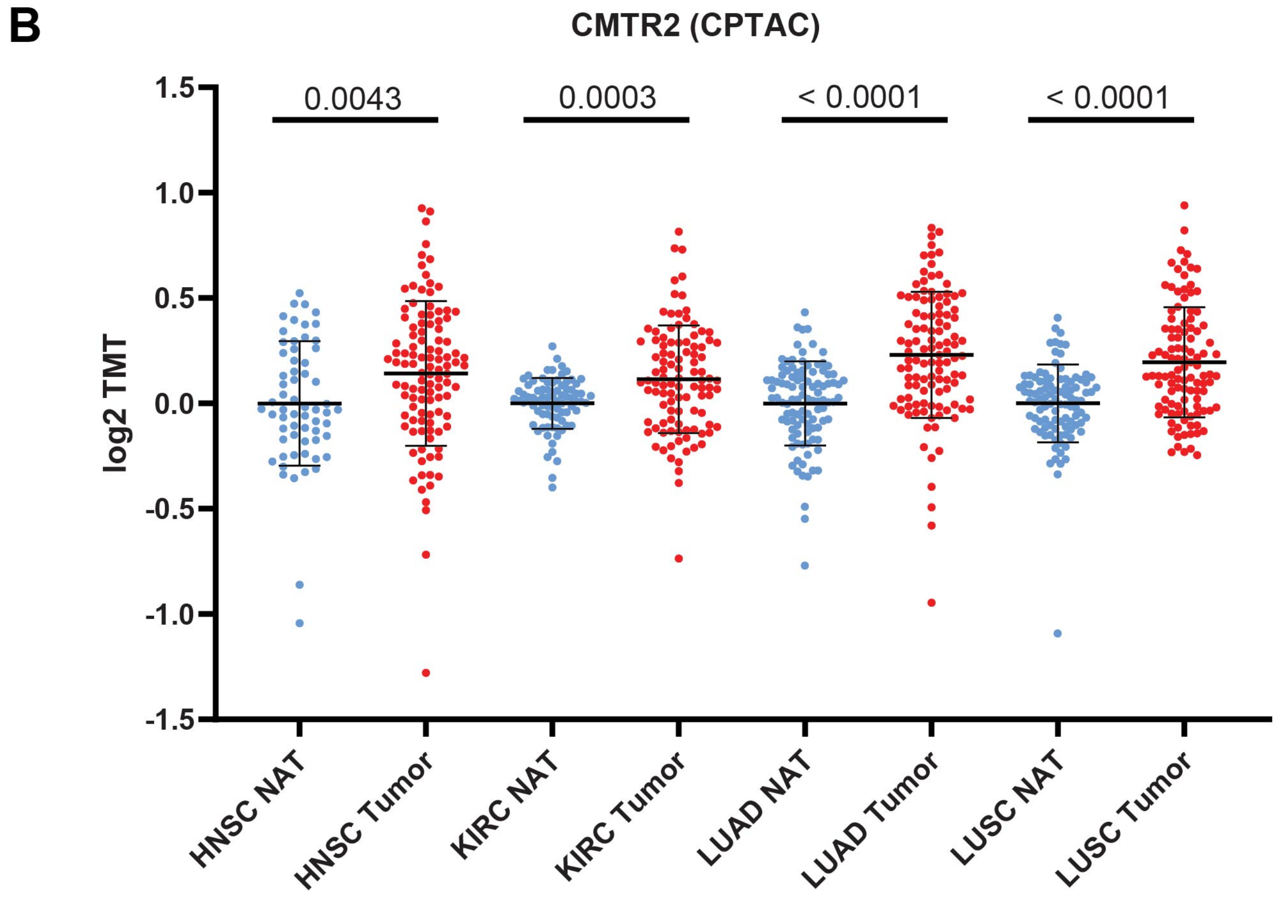

### Supplementary Figure 2

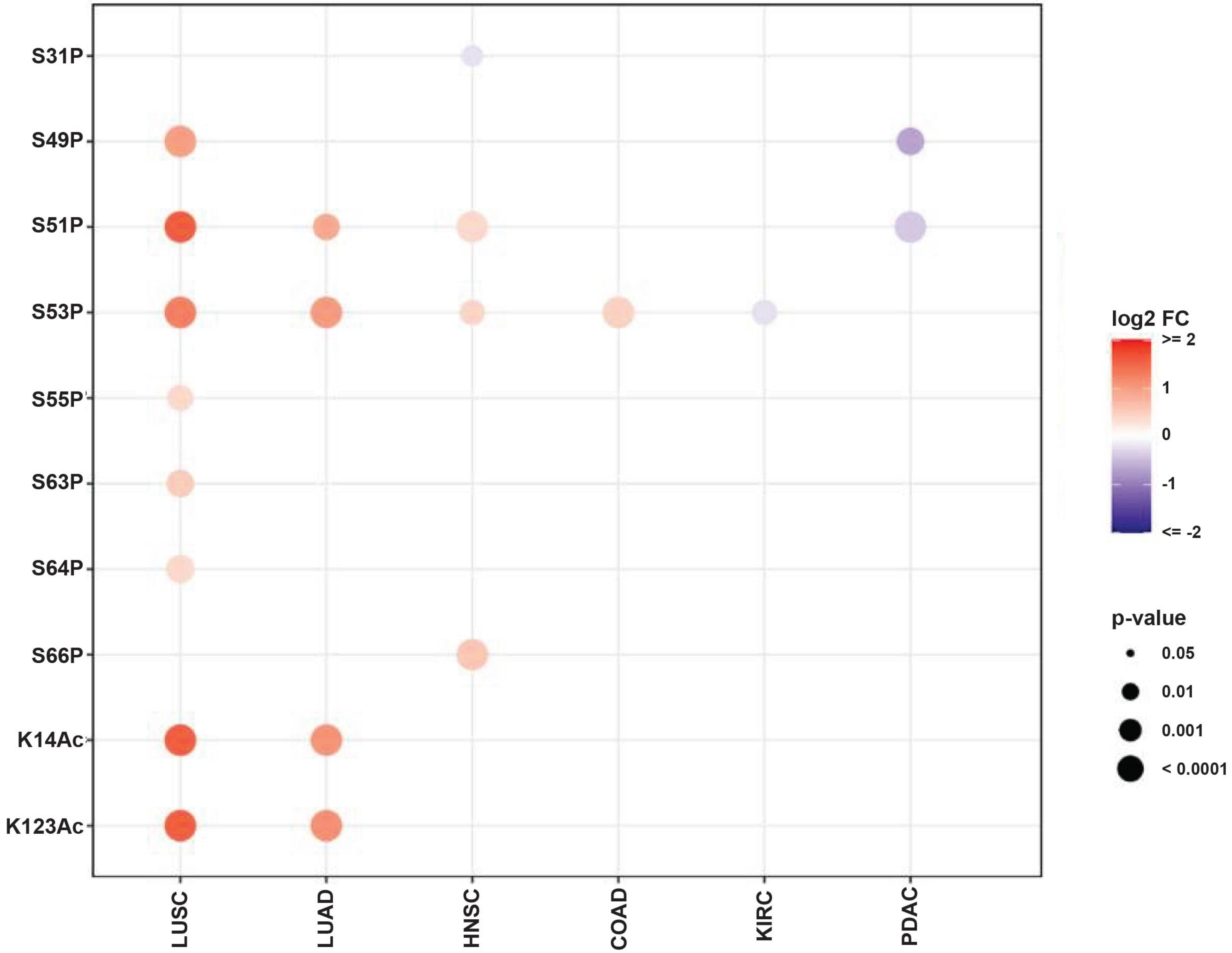

### Supplementary Figure 3

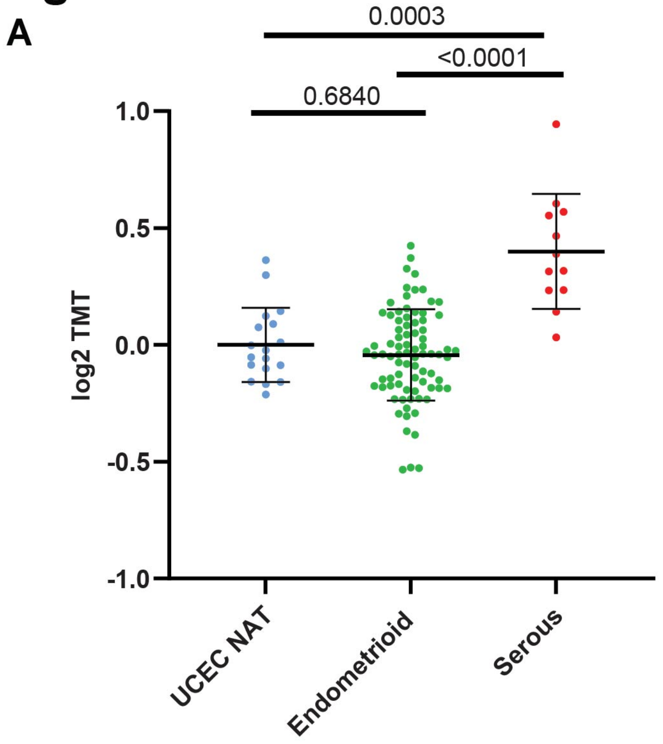


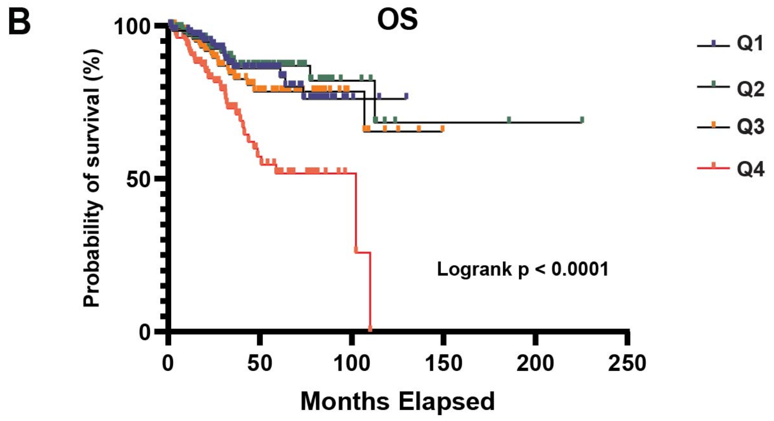


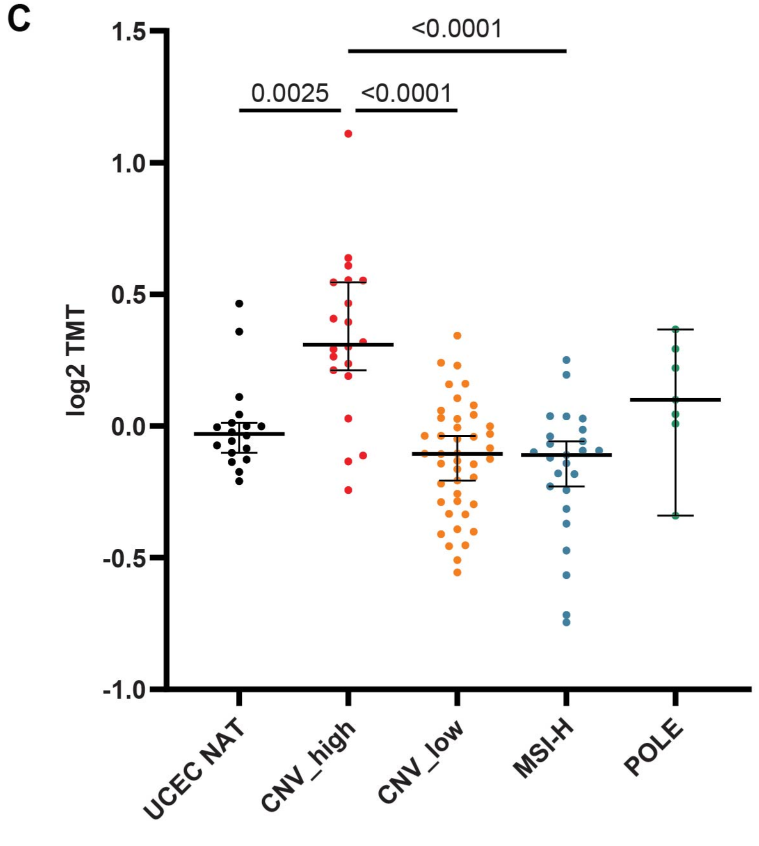

### Supplementary Figure 4

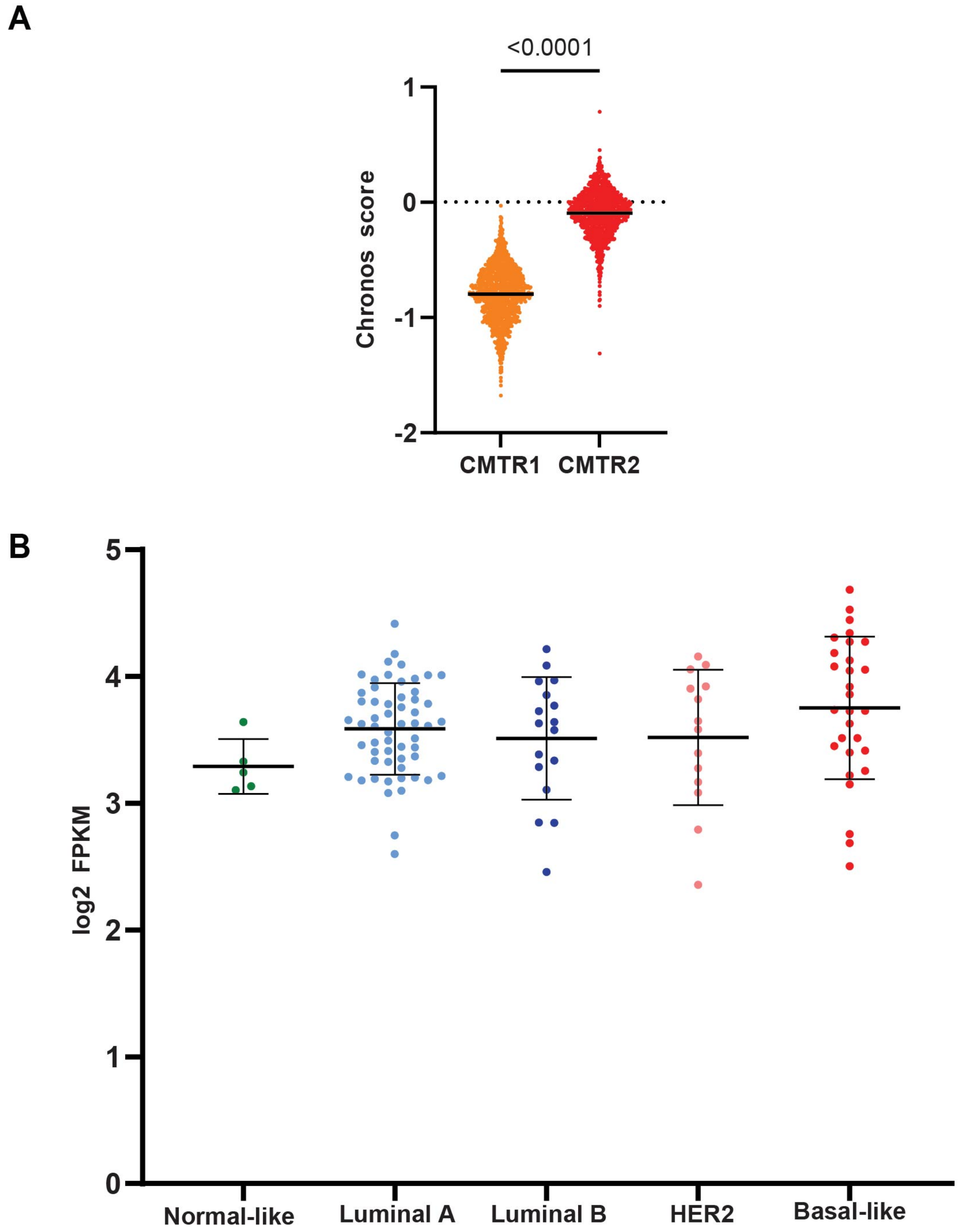

### Supplementary Figure 5

A

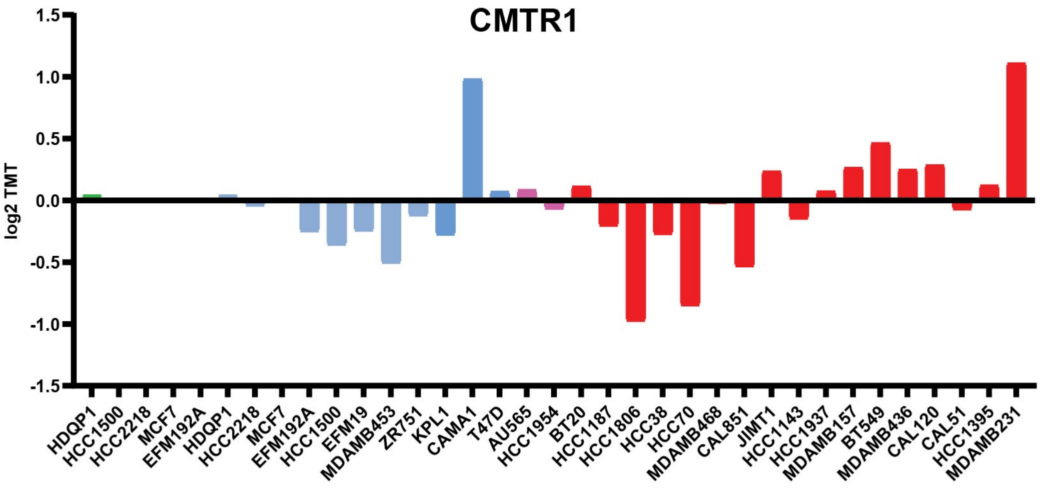


B

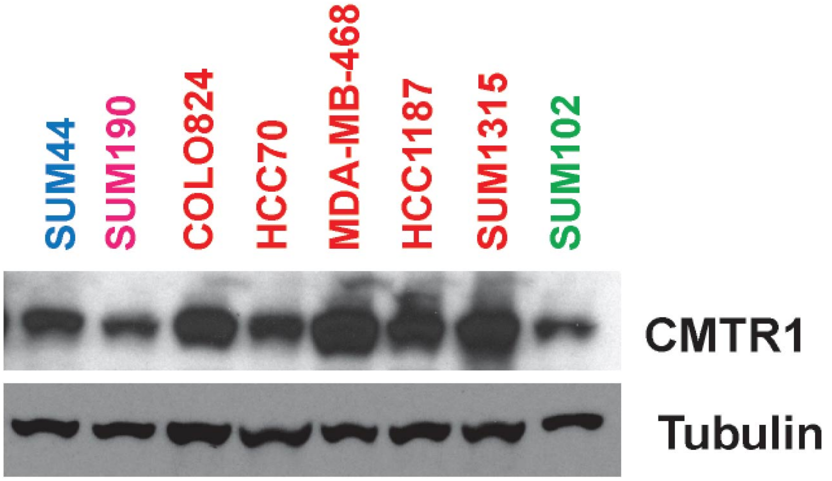

**Supplementary Figure 6**

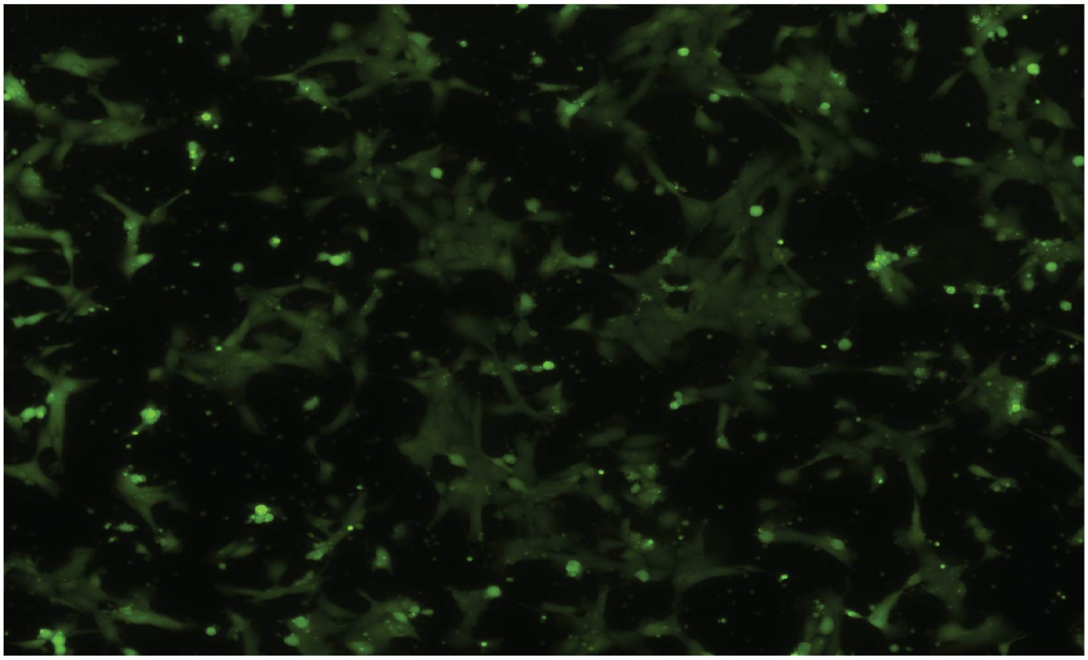

4T1-WT

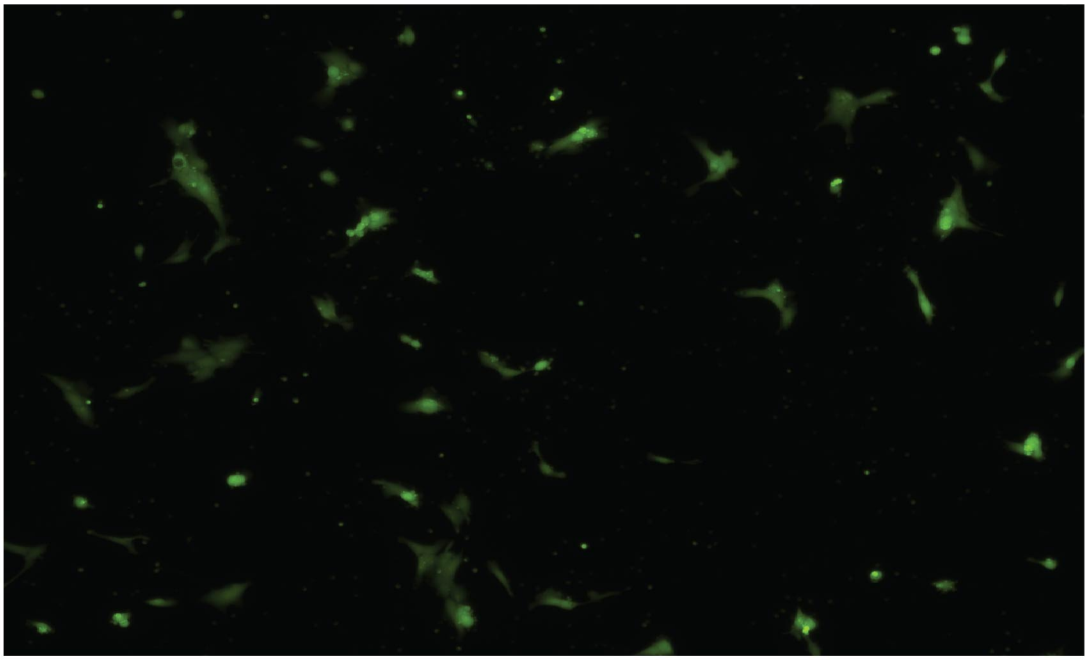

4T1-CMTR1-KO

#### Supplementary Figure 7

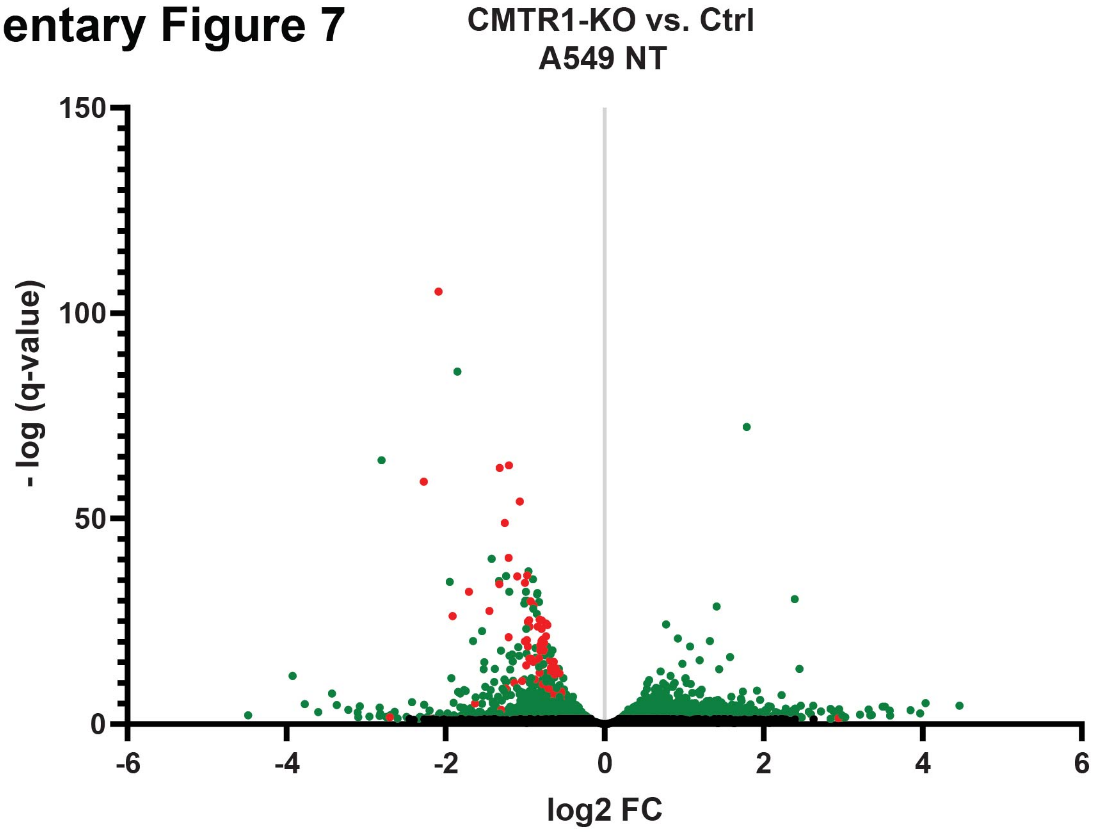


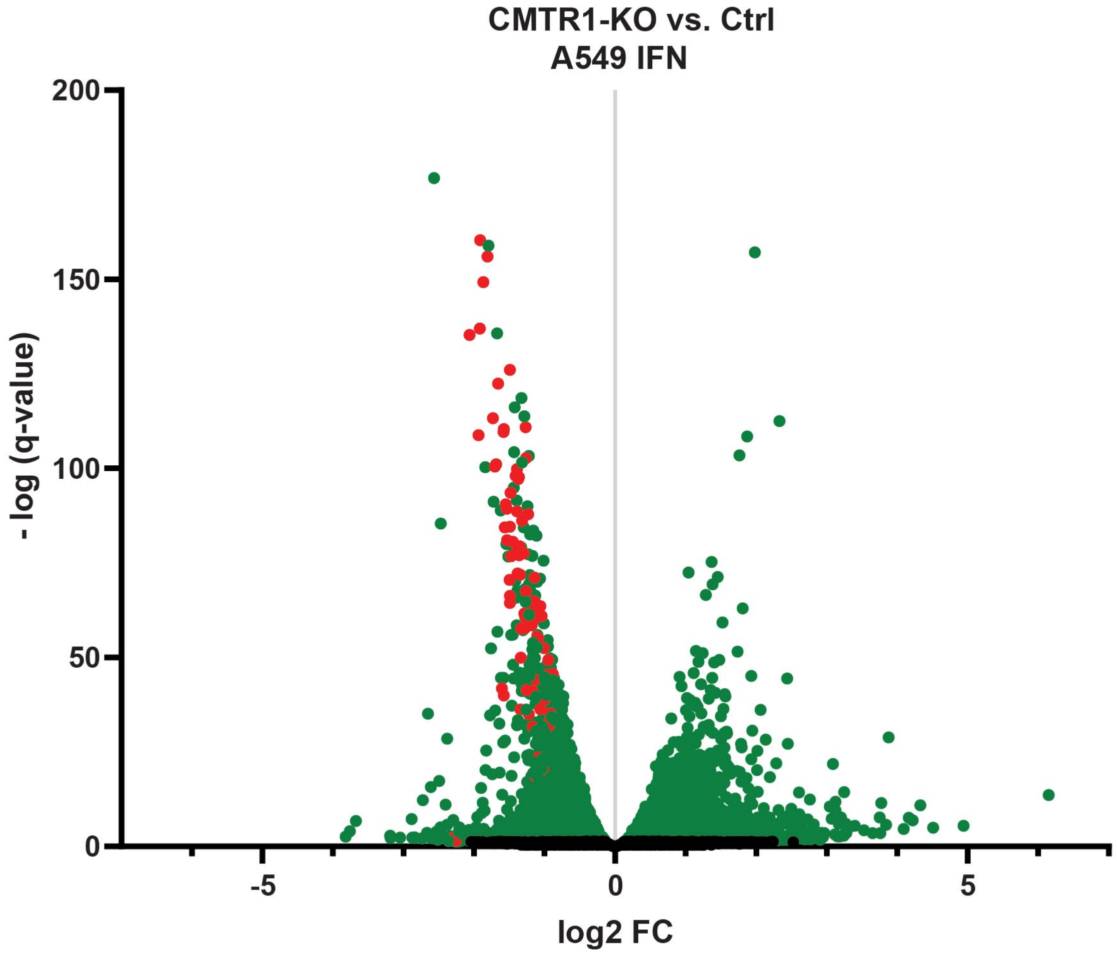

### Supplementary Figure 8

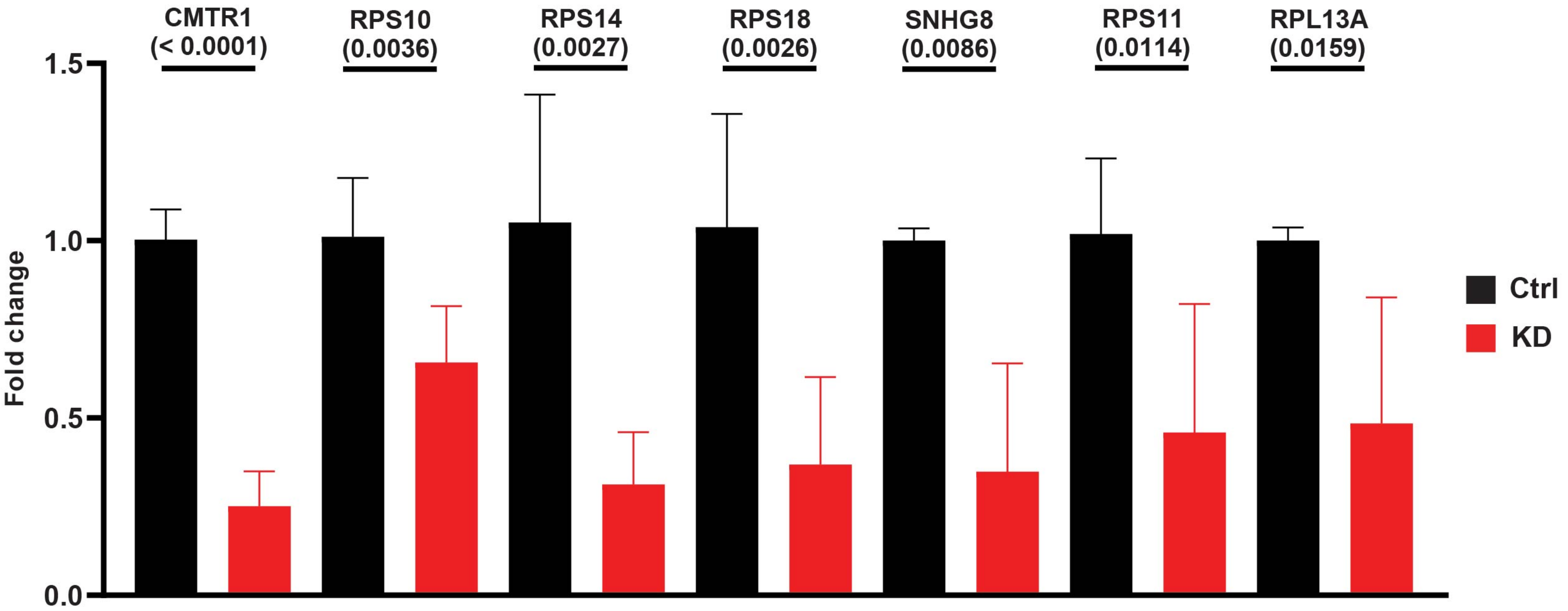

**Supplementary Figure 9**

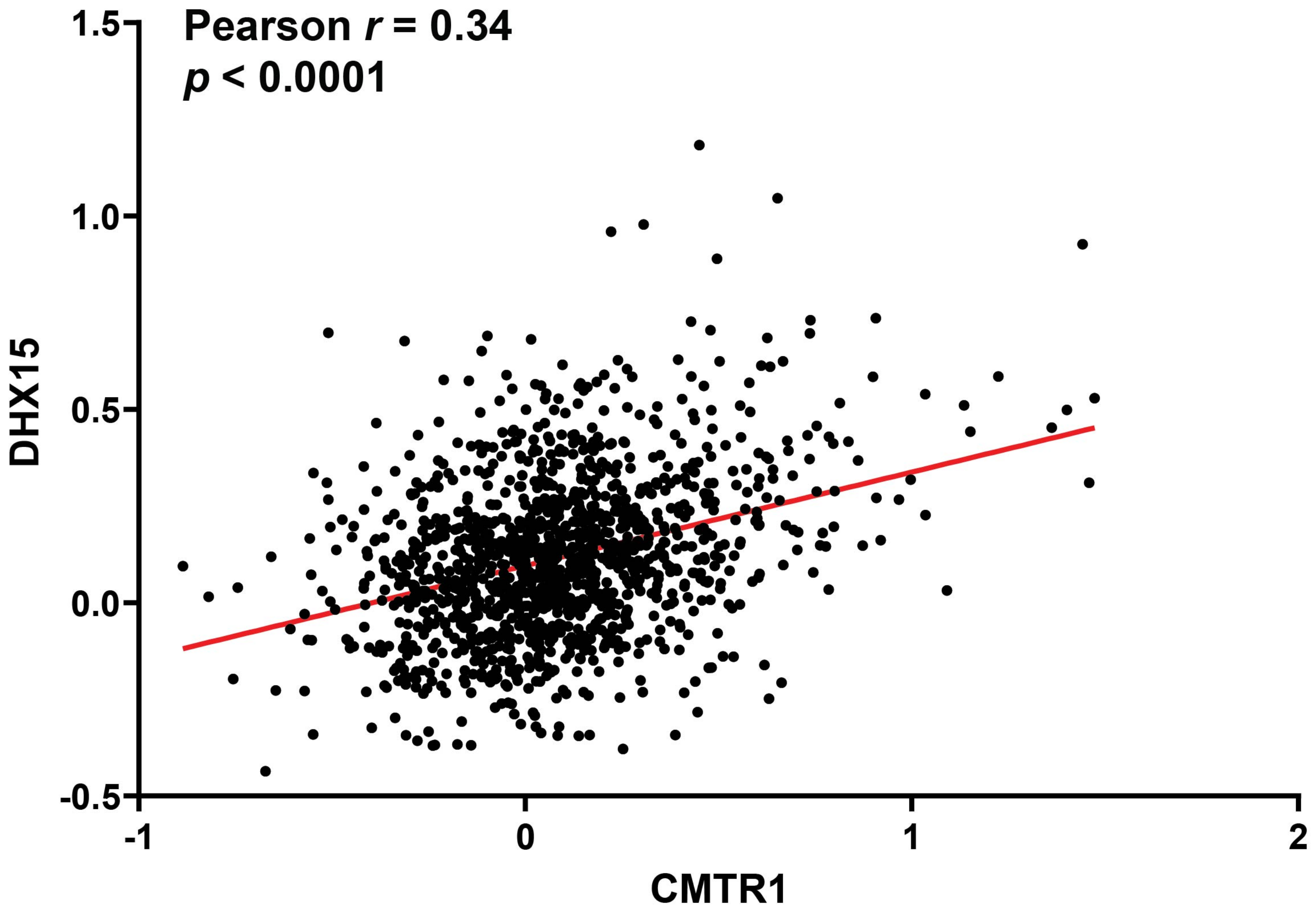
